# On-slide Preparation of *Caenorhabditis elegans* Towards Quantitative, High-Resolution LA-ICP-TOF Mass Spectrometry Imaging

**DOI:** 10.64898/2026.01.08.698490

**Authors:** Aidan J. Reynolds, Aaron Sue, Keith MacRenaris, Thomas V. O’Halloran, Tian (Autumn) Qiu

## Abstract

Metal homeostasis is a complex process wherein essential metals serving structural, catalytic and regulatory roles are acquired, trafficked, and exported once they are present in excess. Understanding changes in metal content and localization in heterogenous tissue types is critical to understanding fundamental physiology as well as a growing number of disease states. Laser ablation inductively coupled plasma time-of-flight mass spectrometry (LA-ICP-TOF-MS) imaging is a powerful technique for untargeted quantitation and mapping of metals in biological systems. While the nematode *Caenorhabditis elegans* (*C. elegans*) is a well-established model organism for fundamental biological research and metal-based diseases, there have been few reports of mass spectrometry-based imaging of *C. elegans*, mostly due to challenges preparing samples that maintain the native distribution of the elements. In this study, we developed an embedding, quantitation and imaging workflow that preserves *C. elegans* using 3D-printed uniform layer media application tools (ULMATs). Multiple embedding media were evaluated, and petrolatum, commercially known as Vaseline™, stood out for its performance in preserving *C. elegans* for imaging applications. Worms were subjected to microscopy and LA-ICP-TOF-MS imaging where we achieved a 2-μm spatial resolution by over-sampling laser shots during ablation. Quantitative elemental maps were obtained using a series of gelatin standards that were sectioned at a 40-μm thickness to closely mimic the average tissue ablation depth of a Day 1 gravid adult *C. elegans*. Our results establish a new workflow for comprehensive elemental profiling of *C. elegans* using LA-ICP-TOF-MS, which holds high potential for future spatial metal biology research with *C. elegans*.

**TOC:** 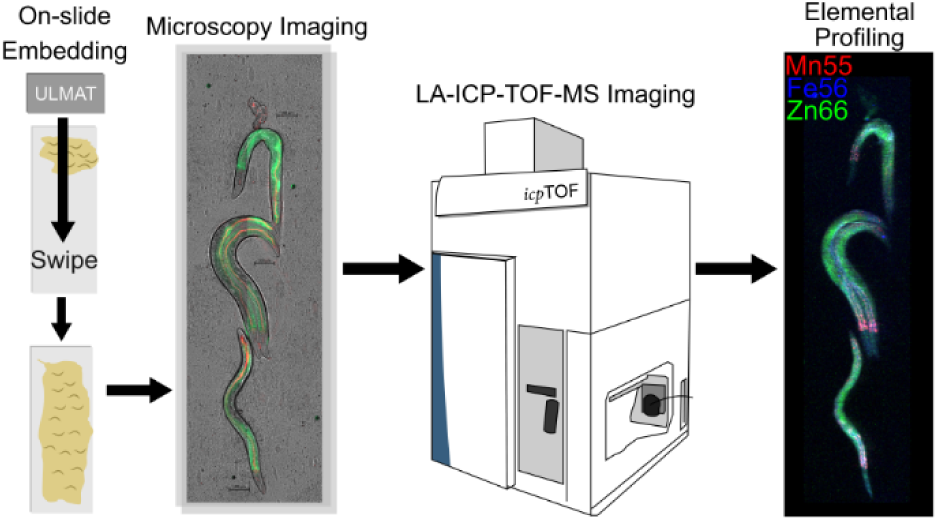

## Introduction

Acquiring, trafficking, and localization of metals in biological systems is essential for physiological processes. Metals can act as structural or catalytic cofactors in cellular pathways such as gene transcription, cell proliferation and enzymatic reactions.^1,2^ Endogenous metal homeostasis can be altered due to non-essential metal exposure in the environment,^3^ pathogenic infection,^4,5^ autoimmune disorders,^6^ and diseases such as Wilson’s disease and hemochromatosis.^7,8^ Tissues possess multiple cell types resulting in highly heterogeneous systems within an organism. This heterogeneity results in the discrete localization of metal ions, metalloproteins, and metal-binding ligands with respect to individual cell types and their unique biological functions, producing an inorganic phenotype. Excessive exposure to non-essential metals in the environment can disrupt endogenous metal homeostasis and changes in the subcellular distribution and activity of biologically relevant metals.^9^ Alterations in the spatial distribution of metals such as iron in tissues can result in oxidative stress leading to cell death.^10^ Understanding the spatial distribution of metals provides a critical dimension of characterization by relating cell types co-localized with metals *in situ* when investigating metal-related biological processes, and perturbations in metal biology within diseases or disease-states.^11,12^ The importance of contextualizing the spatial distribution of elements in biological systems has resulted in the development and advancement of elemental imaging techniques in recent years to support the growing field of spatial metallomics.^13–16^

A collection of techniques is available to map the spatial distribution of metals in biological samples, including X-ray fluorescence microscopy (XFM) methods, metal-binding fluorescent probes, and mass spectrometry imaging (MSI).^17–19^ While these techniques all seek to quantify and visualize the distribution of elements, each offer advantages and disadvantages. X-ray fluorescence microscopy techniques are non-destructive and offer high spatial resolution imaging (100 nm – 50 μm) with moderate sensitivity and can image large areas of tissue simultaneously, however, XFM struggles with light metal ion imaging.^17^ Furthermore, synchrotron-XFM instrumentation is not readily accessible due to high cost and limited facilities. Metal-binding fluorescent probes serve as another non-destructive metal imaging technique, where fluorophores conjugated with metal chelators changes structure upon metal binding and produce optical signals.^20^ While metal-binding fluorescent probes offer good spatial resolution and offers possibilities for live cell imaging, probes can only bind to free metal ions. Additionally, these probes can be promiscuous and bind to unintended metal ions or adventitious sites which affect accuracy of metal imaging. Finally, laser ablation inductively coupled plasma mass spectrometry (LA-ICP-MS) imaging offers moderate spatial resolution (5 – 100 μm) with excellent sensitivity, can image large areas of tissue, and is capable of imaging light metal ions.^19^ When coupled with time-of-flight (TOF) mass analyzers, LA-ICP-TOF-MS offers simultaneous detection of most elements on the periodic table in one scan.^21^ While the destructive nature of LA-ICP-MS imaging should be noted, its ability to perform simultaneous detection of multiple elements while balancing detection specificity, sensitivity and spatial resolution, makes it an ideal choice for spatial metallomics.^18^

Many model organisms have been used for the study of metal biology, with the majority of spatial metallomics done on *Daphnia magna,* zebrafish, and mice.^17^ The nematode *Caenorhabditis elegans* (*C. elegans*) is an emerging model organism for metal biology research, offering advantages such as its conserved genetic homology with mammals (60%-80%), tractable genetics, characterizable phenotypes, and ease of culture.^22–24^ *C. elegans* is approximately 1 mm in length at the adult hermaphrodite stage, has distinct tissues and cell types, reproduces rapidly within a short developmental period (approximately 3 days), and possesses many shared mechanisms of metal transport and homeostasis as mammals.^24–26^ Despite the promise of *C. elegans* in metal biology research, few studies have shown high-spatial resolution metal distributions using LA-ICP-MS, likely due to challenges in sample preparation.^27^ An ideal sample preparation method for metal imaging needs to both preserve the worm’s anatomical structures and avoid delocalizing metals. Previous studies relied on sample preparation steps by intentional desiccation or fixation, where desiccation resulted in significant tissue damage.^28–31^ Fixation techniques preserve the worm’s anatomy but can differentially impact metals, thus affecting the localization of metals and introducing interference in detection, particularly for LA-ICP-MS.^32^ Therefore, a method that can preserve the anatomical structure of *C. elegans*—a sensitive organism that is prone to desiccation and tissue damage during handling—and avoid aggressive use of chemicals for sample processing is highly desired for spatial metallomic studies using *C. elegans*. In addition, due to the small size of *C. elegans*, high spatial resolution, ideally at cellular levels, is desired to visualize metals in specific tissues for functional interpretations.

In this study, we established a novel sample preparation protocol for whole-animal elemental imaging of *C. elegans* by LA-ICP-TOF-MS using petrolatum as an embedding media and achieved 2-μm spatial resolution with laser ablation oversampling. Petrolatum showed excellent preservation of *C. elegans* while retaining sufficient optical transparency for microscopy visualization, possessed the necessary nonvolatility for long-term storage, and was elementally pure allowing for high quality LA-ICP-MS imaging. With the assistance of a home-designed 3D-printed tool, uniform layer media application tool (ULMAT), petrolatum was applied as a uniform layer on top of charged microscope slides for on-slide embedding, freezing, and subsequent elemental analysis of *C. elegans*. Using an eyelash pick, an optional and additional step was also used to extract deceased worms from the embedding media layer for elemental imaging with minimal embedding media matrix effects. Microscopic analysis showed excellent preservation of worm tissue after sample preparation with petrolatum and liquid nitrogen freezing. Through several time-series experiments at ambient conditions, we evaluated sample integrity under different sample preparation approaches to determine optimal imaging time frames such that tissue decay and metal delocalization did not impede the accuracy of elemental imaging. We then characterized LA-ICP-TOF-MS imaging artifacts indicative of the stages of sample desiccation. We applied an over-sampling technique in LA-ICP-TOF-MS imaging and achieved a 2-μm spatial resolution, providing high resolution imaging of *C. elegans* elemental content. Finally, using metal-spiked gelatin standards and ImageJ-based data analysis pipeline, we developed quantitative LA-ICP-TOF-MS imaging workflow and successfully produced quantitative elemental profiling of key biological elements in *C. elegans*.

## Experimental Section

### Materials and Reagents

Optimal cutting temperature compound (OCT), Difco^TM^ agar, granulated peptone, and sodium hydroxide pellets (98%), and Epredia charged microscope slides were purchased from Fisher Scientific (Hanover Park, IL, U.S.A). Gelatin from cold water fish skin, porcine gelatin, carboxymethyl cellulose (∼90,000 Da), polyethylene glycol solution (1000 Da, ≥99%) (50% v/v in H_2_O), Glycerol (≥99%), sodium chloride (≥99%), calcium chloride dihydrate (≥99%), potassium phosphate monobasic (≥99%), potassium phosphate dibasic (≥99%), magnesium sulfate (≥99%), cholesterol (≥99%), and sodium phosphate dibasic (≥99%), and sodium phosphate monobasic (≥99%) were purchased from Sigma-Aldrich (St. Louis, MO, U.S.A). Concentrated clorox germicidal bleach (8.25% sodium hypochlorite) was purchased from the Michigan State University Stores. Indium tin oxide (ITO) slides (70 – 100 Ω) (Delta Technologies, Loveland, Colorado). Vaseline™ petroleum jelly (petrolatum) was purchased over the counter. Anycubic Photon Mono 4 and standard resin for 3D printing was purchased from the Anycubic online store. A custom standard IV-74434: 1000 µg/mL (As, B, Ca, Cd, Co, Cr, Cu, Fe, K, Mg, Mn, Mo, Na, Ni, Pb, Se, V, Zn) was purchased from Inorganic Ventures (Christiansburg, VA, U.S.A).

### Caenorhabditis elegans strain handling

The following *Caenorhabditis elegans* (*C. elegans*) strains were used: N2 (wildtype), and SJ4144 (*zcIs18 [ges-1::GFP(cyt)]*) which expresses cytosolic GFP in the intestine. SJ4144 were cultured on nematode growth media (NGM) plates at 20°C fed with *Escherichia coli* (*E. coli)* OP50-tdTomato strain for fluorescence visualization whereas N2 *C. elegans* were cultured on *E. coli* OP50. These strains were purchased from the Caenorhabditis Genetics Center (Minneapolis, MN, U.S.A). To generate large populations of worms for imaging, synchronized SJ4144 *C. elegans* populations were created by isolating embryos from gravid adults using a bleaching solution (0.5M NaOH, 1.2% NaOCl, 70% H_2_O). Isolated embryos were rinsed with M9 buffer to remove bacteria and incubated overnight.^33^ Once hatched, L1 larvae were transferred to NGM plates seeded with their respective *E. coli* food, and experiments were performed using day 1 gravid adult worms. To minimize stress and potential metallomic phenotype variation in the synchronized F1 generation from bleach synchronization, wildtype *C. elegans* used in quantitative LA-ICP-TOF-MS imaging were synchronized using an egg lay synchronization where 10-20 gravid adults were transferred to seeded NGM plates and allowed to lay eggs in a 3-hour period. Gravid adults were removed after the synchronization period, and the remaining eggs were allowed to develop until day 1 gravid adulthood for imaging.

### 3D-printed tool assisted C. elegans sample preparation for elemental imaging

Day 1 gravid adult *C. elegans* were harvested from the surface of NGM plates by gently washing with M9 buffer and recollecting the worm-containing M9 solution in a microcentrifuge tube. Worms collected this way were centrifuged at 300 g for 1 minute, and the supernatant was removed. Fresh M9 buffer was added to the worm pellet, and the rinsing/centrifugation process was repeated at least 3 times or until the supernatant was no longer turbid due to excess bacterial biomass. The remaining pellet was transferred onto an unseeded NGM plate to allow for excess liquid absorption into the NGM agar for 10 minutes. Once excess liquid was absorbed and worms could freely move on the NGM plate, a plastic spatula was coated in petrolatum and was repeatedly touched across the surface of the NGM plate to pick up 50-100 *C. elegans*. Once the desired number of *C. elegans* were adhered to the petrolatum on the spatula, the petrolatum was transferred to a microscope slide. A 3D-printed tool made using standard resin named the uniform layer media application tool (ULMAT) was used to smooth the petrolatum-transferred worms on the microscope slide, creating a uniform ∼100-μm layer of petrolatum with worms embedded. This tool was printed using an Anycubic Photon Mono 4 resin 3D printer. Essential 3D printing parameters for ULMAT production are listed in **Table S1**. ULMATs were also used to test *C. elegans* embedding with a collection of other embedding media. Embedded *C. elegans* were euthanized by flash-freezing using liquid nitrogen vapors for 3 minutes and then stored at −80°C until microscopy and LA-ICP-TOF-MS imaging. Worms embedded in petrolatum and frozen this way were placed under vacuum to dry for 5 minutes to allow water condensate to evaporate from the surface of the embedding media prior to microscopy and LA-ICP-TOF-MS imaging. Petrolatum-coated worms were produced from petrolatum-embedded worms by removing embedded worms from petrolatum with an eyelash pick and transferring to an exposed surface of the slide. These worms were considered “petrolatum-coated” as a small amount of petrolatum remains on the surface of the worm to keep them hydrated. All sample preparation process is illustrated in **Figure 1A-E**. Brightfield, GFP and tdTomato fluorescence imaging of embedded and coated worms were taken using a Zeiss Axio Imager.Z2 upright fluorescence microscope. Embedded and coated worms were then taken for elemental analysis by LA-ICP-TOF-MS imaging using a TofWerk icpTOF S2. Essential microscopy and LA-ICP-TOF-MS parameters for *C. elegans* imaging are shown in **Table S2** and **Table S3**.

**Figure 1:**
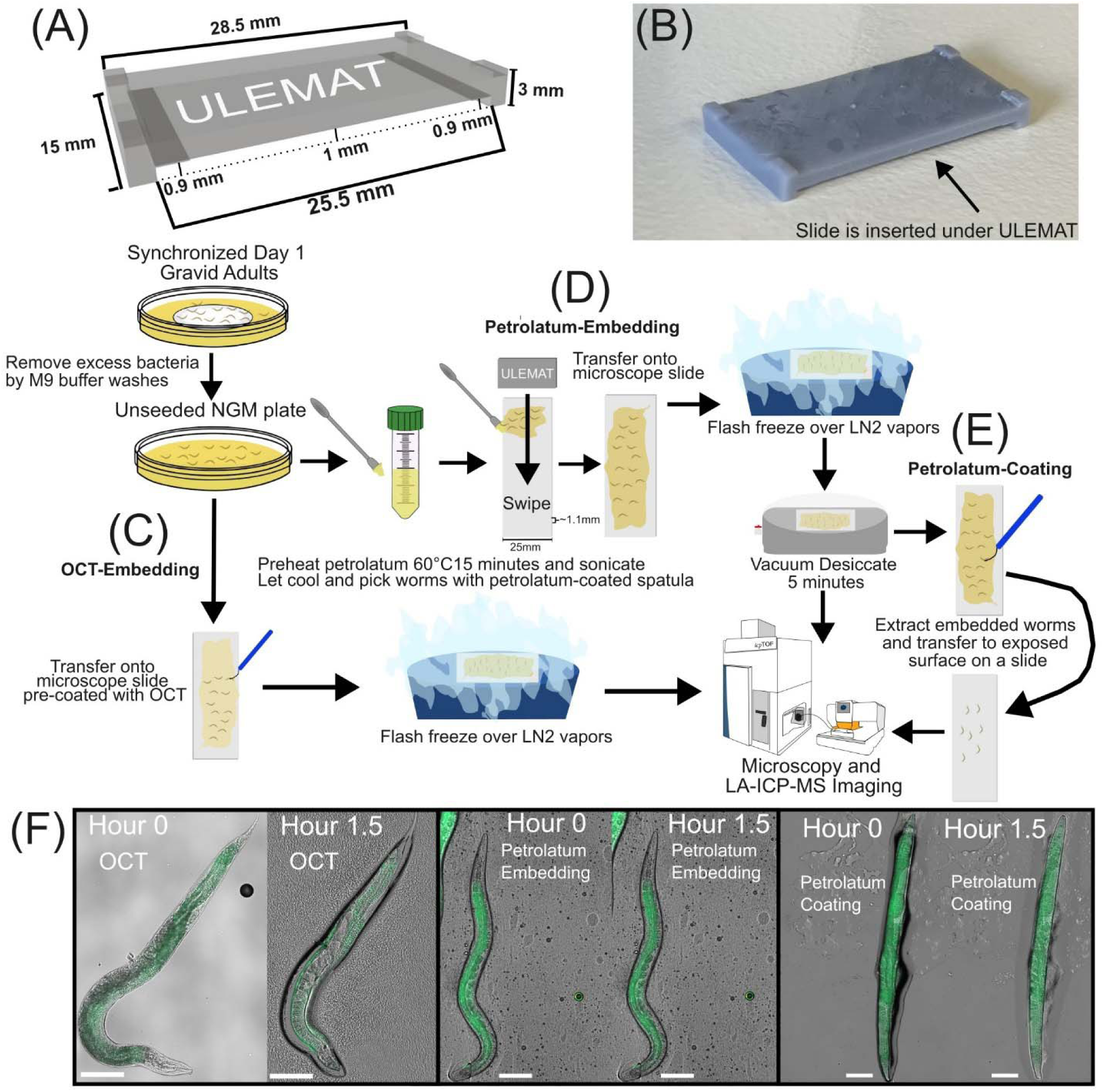
Summary of *C. elegans* sample preparation workflow and short-term visualization of processed *C. elegans* from different techniques. (A) A partially transparent rendering of the uniform layer media application tool (ULMAT) used for embedding and orienting *C. elegans*, (B) and the final 3D printed product tool. (C) Combined *C. elegans* embedding workflow for OCT-embedding, (D) petrolatum-embedding, (E) and petrolatum-coating. A synchronized population of day 1 gravid adult *C. elegans* were rinsed with M9 and transferred to an unseeded NGM plate to let the M9 buffer absorb into nematode growth media prior to the embedding workflows. (F) Appearance of day 1 gravid adult SJ4144 *C. elegans* embedded in OCT, embedded in petrolatum, and coated in petrolatum observed during a 1.5-hour time period visualizing differences in brightfield, GFP fluorescence imaging, and preservation effectiveness. Brightfield black contrast was set to 0 for all microscopy images. OCT-embedded and petrolatum-coated brightfield white contrast was set to 16000 and 9000, respectively. Petrolatum-coated brightfield white contrast was set to 6000. Green fluorescence black contrast was set to 150 and white contrast was set to 1500 for all microscopy images. White scale bars in microscopy images represent 100 μm.

### Monitoring C. elegans preservation post-embedding over time

We first performed monitoring of the same embedded worms over time under a microscope. Petrolatum-embedded worms were frozen by liquid nitrogen vapor for 3 minutes, thawed, and desiccated at room temperature for 5 minutes to remove excess condensate from the slide. Post-processing worms were individually imaged and monitored at the following times for brightfield, GFP, and tdTomato microscopy imaging: 1 hour, 3 hours, 5 hours, 8 hours, 24 hours, and 48 hours at room temperature. Petrolatum-coated worms were imaged at the following times for brightfield and GFP imaging: 1 hour, 3 hours, 5 hours and 8 hours at room temperature. These experiments were repeated 3 times using different synchronized populations of SJ4144 to account for differences in preservation quality due to variations in sample preparation and population.

### Investigating desiccation effects on metal localization in OCT-embedded C. elegans

Day 1 gravid SJ4144 *C. elegans* were embedded in OCT and immediately imaged without the use of gelatin standards. One slide containing SJ4144 *C. elegans* were removed from −80°C storage and underwent microscopy imaging and LA-ICP-TOF-MS imaging within 1 hour at ambient conditions. A second slide from the same population of SJ4144 *C. elegans* were allowed to dry for 1.5 hours prior to microscopy imaging and LA-ICP-TOF-MS imaging to compare distribution of elements within imaged worms. Worms were imaged 2-4 hours post-processing this way. Metal distribution was characterized using ^55^Mn as it exhibited the most discrete localization in *C. elegans*, and its subsequent delocalization would be a strong indicator for morphological changes.

### Elemental quantification of C. elegans

To quantify elemental content of wildtype day 1 gravid adult worm samples, 10% (w/w) porcine gelatin standards doped with either a liquid standard mixture (Inorganic Venture custom standard IV-74434) or sodium phosphate monobasic were prepared at a 40 µm thickness. *C. elegans* and standards were scanned at 5-μm spot size and 2-μm raster spacing with a 3-μm overlap to achieve pseudo-2-μm lateral resolution. To eliminate the initial gelatin standard scan which uses the full 5-μm spot size, the imaging run was interrupted after the first line of each gelatin standard was ablated. The initial lines were removed from the defined ROIs such that the re-queued imaging run would resume ablating with an effective 2-μm spot size. An aliquot of each gelatin standard was measured using an ICP-MS (Agilent 8900 ICP triple quadrupole) to confirm the concentration of each element in each gelatin standard **(Table S4)**. Standards were scanned at equal intervals 10 times for each run, including at the start and end of the scan. A gas blank was collected for 5 seconds before each standard scan. Standard concentrations: 0, 12.5, 25, and 50 ppm (IV-74434) and 1000, 2000, and 4000 ppm (sodium phosphate monobasic). All reported ppm concentrations were produced from w/w dilutions and were assumed to be equivalent to μg/g values. Synchronized populations of N2 *C. elegans* were cultured on *E. coli* OP50 and embedded in OCT, embedded in petrolatum, or coated in petrolatum for initial microscopy imaging, followed by elemental imaging up to 4-hour post-processing. Ten worms from each population were sampled to compare elemental quantitation results and assess embedding media matrix effects. These experiments were performed with new gelatin standard sections each day to assess consistency of quantitation across multiple days of imaging.

### Quantitative metal imaging analysis

Worm dimensions were measured using ImageJ (v1.54p) with the installed macro InteredgeDistance_v1.1_ImageJMacro. Calibrated images were imported into ImageJ and parallel lines were defined along the body length of *C. elegans* images from the nose to the tail tip. Once lines were defined, the macro randomly sampled between 10 and 50 diameters across the body of the worm between the user-defined lines. The average, standard deviation, number of sampled diameters, minimum measured diameter, and maximum measured diameter were exported into a .csv file for further analysis. The non-uniform rounded shape of *C. elegans* was normalized to a cylinder from the average diameter calculated using ImageJ, with the measured worm length serving as the height of the cylinder. The average depth of tissue ablated by the laser was approximated by redistributing the volume of the cylinder to a cuboid of the same length, where the diameter was retained as the width of the cuboid face. The average depth of tissue ablated across the worm’s body was calculated by dividing the volume of the cuboid by the retained diameter and length **(Figure S1)**. The average depth of tissue ablated was used to determine the appropriate gelatin thickness to best reflect the sample type during quantitative imaging and total metal content in worms.

All LA-ICP-TOF-MS images of *C. elegans* were analyzed and rendered in the Iolite software package.^34^ SJ4144 *C. elegans* images were acquired without the use of gelatin standards for investigations into elemental localization. Generated counts per second (CPS) signals for each element channel were used for elemental map visualization in SJ4144 *C. elegans*. CPS scale bar lower limits were set to 0 and upper limits were set to 99^th^ percentile for ^56^Fe and ^66^Zn whereas ^55^Mn was set to 99.9^th^ percentile. Wildtype *C. elegans* images were acquired with gelatin standards for quantitative imaging. Calibration curves were background-subtracted, and the slopes were forced through the origin (0,0). Limits of detection (LOD) and limits of quantitation (LOQ) in were calculated in Iolite as described by Howell, *et al.*^35^ ^31^P quantitation was normalized to the 4000-ppm calibration standard whereas ^55^Mn, ^56^Fe, ^63^Cu, and ^66^Zn were normalized to the 50-ppm calibration standard. Intensity scale bar lower limits were set to 0 and upper limits were defined for ^55^Mn as 99.9^th^ percentile and 99^th^ percentile for ^31^P, ^56^Fe, ^63^Cu, and ^66^Zn unless otherwise stated. Total metal content in *C. elegans* quantitative imaging was calculated by defining a region of interest (ROI) around *C. elegans* in Iolite and exporting individual pixel concentrations to an excel workbook. Concentrations in each pixel are reported as parts per million (ppm) and the sample density was assumed to be 1 such that ppm was equivalent to μg/cm^3^. All negative concentration values from pixels were adjusted to values of 0 for total metal content determination. The measured μg/cm^3^ were converted to total μg by multiplying the concentration by the area of the pixel and the average tissue depth in the individual worm. The remaining pixel values were summed to determine total metal content in individual *C. elegans* and converted to total pg due to the low metal content in worms. While this approach underestimates the tissue depth at the center of the worm and overestimates near the edges of the worm, this approach only compromises the spatial accuracy of pg per pixel values. The total pg of the summed pixels remains uncompromised allowing for consistent whole-worm metal content calculations. To account for developmental variations in worm size contributing to metal content differences, total metal content was normalized to the volume of the worm.

### Statistics

Potential embedding media matrix effects were assessed using one-way multivariate ANOVA (MANOVA) for ^55^Mn, ^56^Fe, ^63^Cu, and ^66^Zn in worms. Prerequisite assumption tests for MANOVA (Shapiro-wilk test, Box’s M test) and MANOVA calculations were performed using the python package “statsmodels.” Post hoc supervised linear discriminant analysis (LDA) was performed on data to visualize which sample preparation approaches were statistically different from MANOVA analysis. LDA was performed using the python package “scikit-learn” and the samples were split into training (80%, or 22 randomly selected worms) and test (20%, or 6 randomly selected worms) sets. ^31^P content in worms was assessed using one-way ANOVA with Tukey’s HSD post hoc test in GraphPad Prism (v10.6.1) to determine if the washing protocol sufficiently removed the phosphate-containing M9 buffer across all prepared worms.

## Results and Discussion

### Embedding media characterization and on-slide embedding of C. elegans in thin layers achieved using a 3D-printed tool

*C. elegans* as a model organism has shown sensitivity to desiccation, and it is therefore important to create sample preparation conditions that retain moisture in worms. Furthermore, this sample preparation process should simultaneously orient *C. elegans* in a uniform plane to produce consistent high-quality imaging across the whole worm body. To achieve this, we developed a simple 3D-printed tool to enable on-slide embedding of *C. elegans* named the uniform layer media application tool (ULMAT) **(Figure 1A-B)**. With a simple swipe of embedding media on top of a microscope slide, the ULMAT can create a uniform thin layer and simultaneously orient worms in a flat plane for imaging. This tool was developed using resin to produce a smoother surface texture than filament-based printing materials. The ULMAT enabled novel sample preparation workflows for on-slide embedding and orienting of worms for imaging applications **(Figure 1C-E)**. The 3D printed tool was custom designed to produce reproducible layers of embedding media approximately 100 μm in thickness on top of microscope slides, however, the ULMATs height can be readily changed for use on slides with different dimensions. This layering thickness minimizes *C. elegans* out-of-plane orientation as they are typically 50-80 μm in diameter depending on their developmental stage.^36,37^ Additionally, this layering thickness provides tumbling space to minimize the probability of worms receiving blunt trauma from the moving tool while not too thick to prevent complete ablation.

Several methods towards hydrating and preserving *C. elegans* for microscopy and LA-ICP-TOF-MS imaging were explored using water-based embedding media such as OCT **(Figure 1C)**, fish gelatin, carboxymethylcellulose (CMC), and hygroscopic additives such as glycerol and polyethylene glycol. We found that all water-based embedding media showed rapid evaporation which resulted in the desiccation and morphology change of *C. elegans* **(Figure 1F, S2A-B)**. Hygroscopic dopants such as glycerol can be added to improve water retention capabilities of these media, however, these molecules typically function as osmolytes,^38^ or penetrating cryoprotectants. These additives displace water from embedded tissues resulting in even more rapid tissue desiccation. Compared to OCT, fish gelatin and glycerol-doped CMC showed better preservation of reporter fluorescence and worm body, while the wrinkled appearance of the worm cuticle indicated desiccation **(Figure S2A-B)**.

As an analog of paraffin, which is commonly used for histology and immunohistochemistry applications,^39,40^ petrolatum, commercially known as Vaseline™, was tested for embedding *C. elegans* **(Figure 1D)**. Petrolatum is hydrophobic, nontoxic, nonvolatile, optically transparent, commercially available, and malleable for the purposes of embedding at room temperature.^41^ Using the ULMAT tool, we found that petrolatum showed excellent preservation of *C. elegans* morphology **(Figure 1F)**. Additionally, even after moving embedded, post-freezing *C. elegans* from petrolatum and transferring to a new slide, the residual petrolatum coated on *C. elegans’* bodies was able to keep worms hydrated **(Figure 1F)**. Early attempts at directly applying petrolatum from the Vaseline™ container showed bubbles trapped in the media, which could interfere with microscopy and metal imaging **(Figure S3A)**. We showed that air bubbles were effectively reduced by transferring petrolatum to a metal-free conical tube and heating and sonicating at 60°C for 15 minutes to liquify the media, allowing for trapped gases to escape, and then cooling the media for application. Vacuum drying following petrolatum embedding and freezing was also found to effectively remove a majority of bubbles, however, residual bubbles were still present **(Figure S3B)**. We found these residual bubbles showed minimal interference with our imaging analyses.

Additional embedding media preservation characteristics were evaluated through short-term microscopy imaging to characterize embedding media non-specific fluorescence, transparency and preservation effectiveness **(Figure 1F)**. OCT-embedding showed the best initial brightfield and fluorescence microscopy quality but suffered from rapid desiccation. Petrolatum embedding and coating methods both showed excellent hydration and preservation of *C. elegans* and could keep worms hydrated up to 48 hours **(Figure S4)**. It was noted that the GFP reporter in the intestinal cells decayed from initial embedding to 8 hours post-processing whereas brightfield imaging showed no significant changes during this time frame **(Figure S4, S5)**. At 24 hours, we observed morphological changes in embedded worms; most notably shriveling of the head which is indicative of worm death and decay, which can be possibly attributed to rigor mortis and intestinal necrosis.^42^ We similarly observed pharyngeal deformations reported by Galimov *et al.* wherein some *C. elegans* showed rounding at the mouth and crumpling of the isthmus resulting in pharyngeal kinking at 24 hours post-processing **(Figure S6A-B)**. At 48 hours, worms experienced severe changes in shape, size, fluorescence, and orientation. We also noticed minimal fluorescence of the tdTomato reporter in *E. coli* OP50 food between 0-24 hours, indicating that our washing protocol is sufficient to remove bacterial content from worms for immediate LA-ICP-TOF-MS imaging.

Through additional microscopy experiments, petrolatum-coated day 1 gravid adult SJ4144 *C. elegans* were found to undergo desiccation more rapidly than petrolatum-embedded worms and remained hydrated up to 8 hours at ambient conditions **(Figure S7)**. Petrolatum coating also experienced significant non-specific fluorescent signals compared to petrolatum embedding. This non-specific fluorescence could not be corrected by adjusting microscopy imaging parameters or green channel fluorescence histogram properties. We suspect that the whole-body non-specific fluorescence is due to the GFP fluorescent light emitted from the intestinal cells undergoing diffuse reflection in the uneven petrolatum coating the surface of the worm.^43,44^ This diffuse reflection of emitted fluorescent light is not as prevalent in the petrolatum-embedded *C. elegans* due to the more uniform texture of the embedding media layer. Therefore, we determined that microscopy evaluation works better with petrolatum-embedded for *C. elegans* with fluorescence reporters. Altogether, our results showed that petrolatum serves as an embedding media to preserve *C. elegans* morphology, anatomy, and fluorescence reporters over the course of 4 hours according to our time-series monitoring, enabling the subsequent evaluation with LA-ICP-MS imaging.

### Optimizing laser parameters for high spatial resolution LA-ICP-TOF-MS imaging and characterizing metal localization

LA-ICP-MS imaging typically use spatial resolutions of 5-50 μm, which provides adequate spatial resolution for larger tissues from model organisms such as rats, mice, zebrafish, and *Daphnia magna*.^45–47^ Due to the limited size of *C. elegans* being ∼1 mm in length and ∼50 μm in diameter, it is essential to optimize laser parameters to maximize signal and spatial resolution during imaging. We developed high-resolution LA-ICP-TOF-MS imaging methods on embedded *C. elegans* by oversampling the ablation laser of 5-μm spot size with a 3-μm over-sampling to achieve an effective 2-μm spot size resolution. This ensured full ablation of the sample by providing sufficient energy from the laser while maintaining a higher resolution. Elemental signatures found in the glass slide such as ^27^Al were referenced to confirm full ablation had occurred with this over-sampling raster pattern **(Figure S8)**. Early imaging attempts with *C. elegans* on ITO slides showed interfering signals from elemental contaminants present in microscope slides. This resulted in interfering signals in glass such as ^56^Fe from iron oxide (Fe_2_O_3_), or excessive ^112^tin producing doubly charged tin ions which serve as isobaric interferences for ^56^Fe **(Figure S8A)**. Subsequent imaging attempts were performed on Epredia charged microscope slides which were found to contain fewer interfering signals upon ablation **(Figure S8B)**. This laser raster pattern was found to provide enough spatial resolution to allow for consistent resolution of the intestinal lumen, allowing for visualization of ^55^Mn between paired intestinal cells which could not be achieved by using a 5-μm laser profile with no over-sampling **(Figure S9A-B)**. Furthermore, the distribution of ^55^Mn appeared to be altered compared to existing synchrotron-XFM literature,^48^ resulting in large paired punctates between intestinal cells in many worms.

To evaluate preservation effectiveness of different embedding techniques on metal distribution, LA-ICP-TOF-MS imaging was performed on prepared *C. elegans* at different time points at ambient conditions **(Figure 2)**. Initial microscopy images of worms prepared by each technique indicate preservation of C. elegans morphology, anatomy and fluorescence at T=0 **(Figure 2A)**. At 60-minute post-processing, OCT-embedded *C. elegans* experienced metal delocalization, likely due to desiccation within 60 minutes of imaging at ambient conditions, which particularly compromised ^55^Mn localization in the intestine of worms **(Figure 2B)**. Petrolatum-embedded and coated *C. elegans* showed no desiccation and subsequently no delocalization of metals **(Figure 2B)**. Occasionally, incomplete ablation was observed due to uneven embedding **(Figure S10)**, but the low frequency of such observation ensured that there were enough worms for acquiring high-quality LA-ICP-TOF-MS images. Picking out worms from petrolatum to generate petrolatum-coated worms further reduced the chance of having unevenly embedded worms and demonstrated clear metal localization **(Figure 2B)**. The reduction in petrolatum content through this method further suggests that too much embedding media present in the sample can impede ion image accuracy and reproducibility. However, without embedding media to serve as a buffer between the laser and microscope slide, excessive ablation of the charged microscope slide face can still introduce interfering signals from glass slide contaminants. While these interfering signals do not impact the accuracy of *C. elegans* imaging, these signals can interfere with elemental quantitation scaling and overall image quality. Transferring petrolatum-coated *C. elegans* to the surface of the petrolatum layer instead of an exposed glass region on the charged microscope slide can possibly remove these imaging blemishes. Artifacts of desiccation were identified using OCT-embedded *C. elegans* to characterize the severity of metal delocalization using ^55^Mn as a benchmark element **(Figure 2C)**. From OCT-embedded high spatial resolution imaging data sets, it was further confirmed that OCT-embedded *C. elegans* experienced morphological changes between microscopy imaging and LA-ICP-TOF-MS imaging, likely due to rigor mortis changing worm orientation or desiccation. ^55^Mn in existing synchrotron-XFM imaging shows discrete localization along the intestinal lumen,^48^ which can be observed in OCT-embedded *C. elegans* that were imaged by LA-ICP-TOF-MS under short imaging times. However, OCT-embedded worms sitting for longer times at ambient conditions post-processing induced gradual delocalization resulting in total delocalization of metals by 3 hours, further emphasizing that water-based embedding media are not suitable for preservation of *C. elegans*.

**Figure 2:**
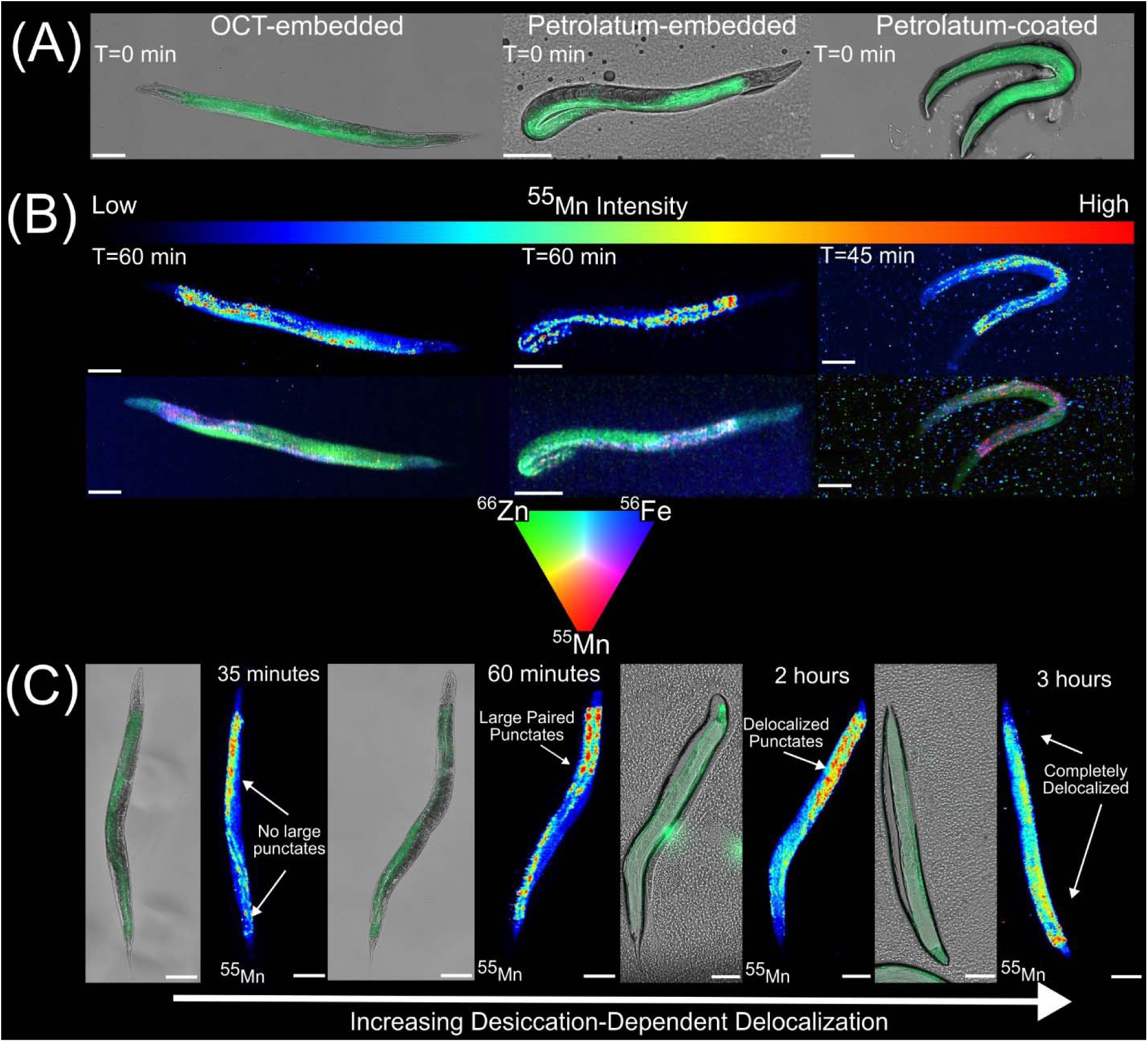
Investigating elemental distribution in *C. elegans* from different sample preparation techniques. (A) Microscopy imaging of day 1 gravid adult SJ4144 *C. elegans* prepared by OCT-embedding, petrolatum-embedding, and petrolatum-coating. (B) Microscopy images were taken shortly after removing samples from −80°C storage and designated as T=0 for ambient condition imaging. LA-ICP-TOF-MS images visualizing ^55^Mn and multi-element overlay of ^55^Mn (red), ^56^Fe (blue), and ^66^Zn (green) (B). LA-ICP-TOF-MS imaging of worms were completed within 60 minutes, with their completion times listed in the corresponding ^55^Mn ion images. (C) Metal delocalization in OCT-embedded *C. elegans* was characterized from hours 0-4 at ambient conditions. Ideal ^55^Mn localization found along the intestinal lumen, with progressive delocalization observed in with respect to time and desiccation progression within the laser ablation stage. ^55^Mn ion intensities were extracted as counts per second, with the lower limit set to 0^th^ percentile and the upper limit set to 99.9^th^ percentile for all ion images. White scale bars in microscopy images and LA-ICP-TOF-MS ion images represent 100 μm.

### High-spatial resolution LA-ICP-TOF-MS provide simultaneous quantitation and mapping of multiple elements on same sample

We further developed a data analysis pipeline to quantify the distribution and total amount of elements from LA-ICP-TOF-MS imaging. Unlike cryo-sectioned animal tissues with the same thickness across tissue, *C. elegans* possesses a tapered, cylindrical body shape with uneven thickness across its length and width, which complicates the quantification from imaging because the laser will ablate different thicknesses across its body. Approximations of *C. elegans*’ dimensions have been reported previously by treating *C. elegans*’ shape as a cylinder.^49^ We further approximate the cylinder to a rectangular cuboid to obtain a uniform thickness by matching the *C. elegans* body volume and length such that standards of adequate thickness could be produced for quantitative imaging **(Figure S1)**. Through ImageJ analysis, we determined the average diameter and length of the body of an early day 1 gravid adult (1-3 hours into adulthood) wildtype *C.* to be 52.12 um, and 933.02 um, respectively **(Figure S11A-B)**. These values generally agree with existing studies on *C. elegans* growth dynamics, ^49,50^ suggesting that the ImageJ macro produces accurate measurements of *C. elegans*.

The average tissue ablation depth across the entire body of *C. elegans* from this cylinder normalization was furthermore found to be 40.93 um **(Figure S11C)**. From these measurements, the volume of *C. elegans* could be calculated for further insight into how metal content correlates with growth dynamics or metal-content normalization **(Figure S11D)**. As gelatin is the most conventional standard material for animal tissue for LA-ICP-TOF-MS imaging,^51,52^ from the calculated average tissue ablation depth, we determined that sectioning gelatin standards doped with element standards at a 40-μm thickness would sufficiently reflect the sample type’s average thickness.

We demonstrated comprehensive elemental quantification and imaging of most elements on the periodic table simultaneously including biologically relevant elements such as ^31^P, ^55^Mn, ^56^Fe, ^63^Cu, and ^66^Zn in *C. elegans* using LA-ICP-TOF-MS technology. **Figure 3** shows an example of a petrolatum-coated worm. Existing quadrupole-based LA-ICP-MS technology is unable to measure all elements simultaneously and is only suitable for imaging a small subset of isotopes. Previous LA-ICP-MS studies on *C. elegans* have only generated elemental imaging results showing one or two elements and their respective isotopes.^29–31^ By overlaying elemental images with brightfield microscope images, we were able to observe discrete visualization of *C. elegans* anatomy for the assignment of elemental localization **(Figure 3)**. ^31^P was found throughout the worm’s body, and most strongly accumulated in the intestinal cells and posterior region of the worm. ^31^P is an essential element in biomolecules such as DNA, phospholipids, energy-storage molecules such as ATP, and as such its abundance can reflect the cell size and activity across regions of tissue.^53^ ^55^Mn exhibited the most discrete localization of imaged elements throughout the intestinal tract of *C. elegans* with a punctate pattern. This defined distribution of ^55^Mn has previously only been visualized using synchrotron-XFM technology and was firstly achieved in this work with LA-ICP-MS imaging.^48^ ^56^Fe showed some increased background noise due to the presence of the isobaric interference ArO^+^, however, we used kinetic energy discrimination within the collision cell and improved ^56^Fe signals. ^56^Fe and ^55^Mn showed higher relative abundances in the anterior intestine in likely the INT1 cells, with significant localization to the intestinal cells throughout the worm body. However, ^56^Fe did not exhibit a punctate distribution like ^55^Mn throughout intestinal cells. While an essential element, ^63^Cu was found in low abundance throughout *C. elegans* potentially due to the organism’s reduced use of the element in biological processes.^54^ Improved quantitative imaging of ^63^Cu can be achieved by using LA-ICP-QQQ-MS imaging if copper biology is of interest. While ^55^Mn and ^56^Fe showed enrichment in *C. elegans*’ anterior intestine, ^66^Zn showed little or no enrichment in the anterior region of *C. elegans* and instead showed increased abundance in the posterior region of imaged worms. Synchrotron-XFM imaging has revealed similar zinc distributions with posterior intestinal cells in Day 5 *C. elegans* possessing high zinc content,^48^ however, our effective 2-μm spatial resolution is not sufficient to resolve the zinc-rich gonad arms from the intestine in *C. elegans.* Additionally, the orientation of *C. elegans* during imaging may confound resolving the gonad arms from the intestine. Multiple elements can be overlaid to generate elemental maps that highlight differences in metal compartmentalization in worms.

**Figure 3:**
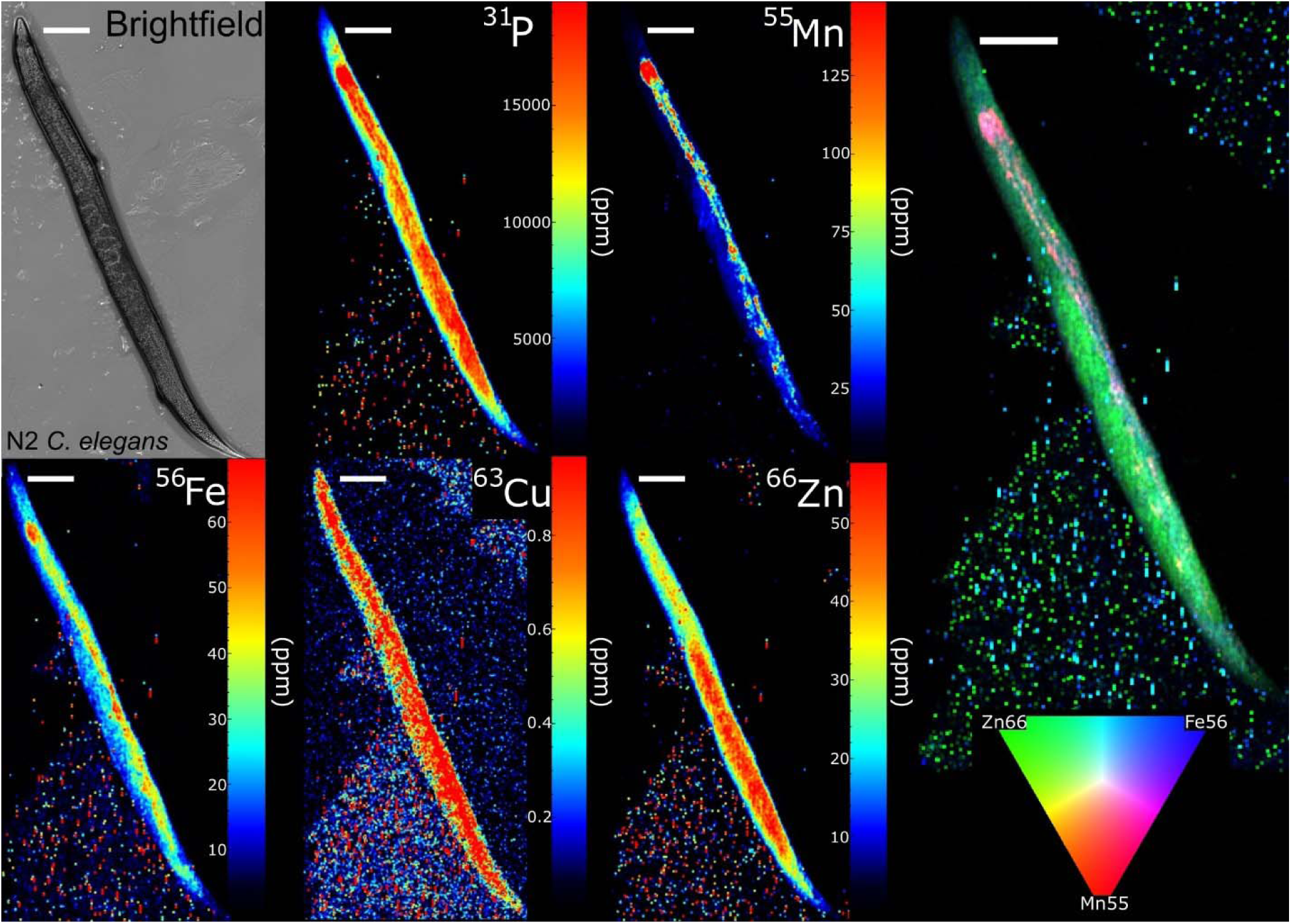
Brightfield microscopy and comprehensive elemental imaging of a petrolatum-coated day 1 gravid adult wildtype N2 *C. elegans*. LA-ICP-TOF-MS imaging was able to perform simultaneous quantitation of ^31^P and essential trace elements including ^55^Mn, ^56^Fe, ^63^Cu, and ^66^Zn, and generate a multi-element overlay image of several endogenous metals. Lower limits for all elements were set to 0 ppm, and upper limits were set to 99^th^ percentile for all elements except ^55^Mn which was set to 99.9^th^ percentile. Brightfield black contrast and white contrast were set to 0, and 9000 respectively. White scale bars represent 100 μm.

Petrolatum-embedded worms were also subjected to quantitative LA-ICP-TOF-MS imaging, which produced similar elemental distributions with respect to anterior versus posterior localization **(Figure S12A-G)**. Some differences in elemental distribution with respect to the intestinal cells were observed with ^31^P and ^56^Fe signals **(Figure S12B, 12D)**. This variation could be attributed to embedding media matrix effects altering plume properties and impacting spatial clarity in imaging or could be due to biological variation. Further elemental imaging must be performed to adequately characterize the spatial metallome of *C. elegans* as worm orientation, embedding media thickness, and developmental variations may contribute to observed differences in metal distribution. In both petrolatum-prepared sample sets, quantitative LA-ICP-TOF-MS imaging of *C. elegans* provided significant coverage of elements for quantitation and visualization with much greater throughput than synchrotron-XFM technologies, allowing for imaging of multiple *C. elegans* in a matter of hours.

### Investigating potential embedding media matrix effects through single worm quantitation

The robustness of quantitative LA-ICP-TOF-MS imaging was investigated through multi-day quantitative imaging experiments for OCT-embedded, petrolatum-embedded and petrolatum-coated *C. elegans* to characterize calibration curve reproducibility and uncover potential embedding media matrix effects between methods. Calibration curves produced from raw data showed significant inter-day variations in sensitivity, however, calibration curves generally produced satisfactory R^2^ values (>0.95) **(Figure S13A-E)**. Investigations into individual imaging runs showed ion intensity and slope sensitivity drift when raw data from gelatin calibration curves were visualized **(Figure S14A-E).** In many experiments, signal intensity and calibration curve slopes decreased in sensitivity as imaging runs progressed. This signal intensity and slope sensitivity drift is due to data acquisition limitations of TOF-MS detectors with single-pulse laser systems, where variations in laser fluency and beam size may change during an imaging run.^55^ Other TOF-MS imaging systems such as matrix-assisted laser desorption/ionization (MALDI) TOF-MS systems typically employ bursts of laser pulses where the spectral output is averaged between all laser shots per pixel which accounts for pulse-to-pulse variations in the laser. Additional factors such as temperature fluctuations may also affect signal acquisition by reducing detector sensitivity, efficiency of vaporization, ablated particle size, and ionization.^56^ Limits of detection (LODs) and limits of quantitation (LOQs) from inter-day imaging experiments showed large relative standard deviation percentages (RSD%) **(Table S5)**. Despite the large variation in instrument performance observed from inter-day imaging, there was no significant impact on elemental quantitation as all LODs and LOQs fell below the lowest gelatin standard concentration (12.5 ppm for ^55^Mn, ^56^Fe, ^63^Cu, ^66^Zn; 1000 ppm for ^31^P), except for one imaging experiment where the LOQ for ^31^P was measured as 1278 ppm **(Table S5)**. Instances of severe instrumental drift can be corrected with external software such as AutoSpect to improve data quality.^57^

Total metal content from quantitative LA-ICP-TOF-MS imaging was determined to assess efficacy of washing off the phosphate-containing M9 buffer, embedding media matrix effects, and how biological variance impacted single-worm quantitation reproducibility. Single worm quantitation has typically reported values of metals in the picogram range,^58^ however, these values without normalization can lead to incorrect data interpretations. *C. elegans* experiences rapid growth and development throughout adulthood,^59^ which can result in size-dependent differences in total metal content. Normalizing to phosphorus or sulfur content is a common approach to improving data comparability between biological replicates but is not suitable for investigating matrix effects between groups.^60^ Matrix effects can unilaterally affect all signals acquired by LA-ICP-TOF-MS, including phosphorus and sulfur such that total metal content may change but the ratio of the analyte ion and normalization ion is unchanged. Ultimately, a normalization strategy to account for biological variation using a feature that is independent of matrix effects is desired to allow for investigations into total metal content differences between sample preparation techniques. To improve comparisons between *C. elegans* from different sample preparation conditions, we opted to normalize total metal content from individual *C. elegans* to the respective volume of the analyzed *C. elegans*. The values acquired from this normalization approach also allowed for simple conversion to molar concentration of elements in *C. elegans*.

Unnormalized total element content, volume-normalized total element content, and molar concentrations of metals from OCT-embedded, petrolatum-embedded, and petrolatum-coated *C. elegans* are shown in **Table 1**. ^31^P is the only element where components of the washing buffer may interfere with reproducibility. Among the 8 repeated experiments for washing and imaging *C. elegans*, only 6 imaged worms from 2 days from the petrolatum-coated group showed significantly elevated ^31^P content **(Figure S15)**. These worms were prepared on high humidity days which prevented adequate evaporation and absorption of the washing buffer into NGM plates from the surface of the worms. When these 2 days are excluded from ^31^P quantitation comparisons, the adjusted ^31^P content produces comparable values to ^31^P from OCT-embedded and petrolatum-embedded groups **(Table 1)**. Future LA-ICP-TOF-MS imaging of *C. elegans* should use phosphate-free buffers to wash worms of bacteria and not perturb ^31^P quantitation. Unnormalized essential transition metal content between sample preparation groups also varied significantly, however, when normalized to respective worm volume these adjusted values resulted in better comparison between groups **(Table 1, Figure S16)**. Elemental content variance generally fell between 20% and 45% percent relative standard deviation in both un-normalized and normalized data sets. To our knowledge, this is the first report in the quantitative metal profile of day 1 gravid adult *C. elegans*, and the variance reported in this data may be confounded by the developmental transition from young adult to early gravid adult. Future research should investigate how *C. elegans’* metallome evolves from early adulthood to later stages of adulthood to better contextualize this data and the observed variance to biological variance.

**Table 1:** Metal Content Per Worm Measured by LA-ICP-TOF-MS.

| Elemental Content per Worm. Values shown as average (RSD%) |  |  |  |  |  |
| --- | --- | --- | --- | --- | --- |
| Sample Preparation | Element | Total Content (pg) | Total Content Normalized to Volume (pg/$\mu\text{m}^3$)<sup>a</sup> | Total Content Normalized to Volume ($\mu\text{mol/L}$) | Replicates |
| OCT-Embedded | <sup>31</sup>P | 9139.6 (15.99%) | $5.1 \times 10^{-3}$ (22.5%) | $1.5 \times 10^5$ (22.5%) | n = 10 |
| | <sup>55</sup>Mn | 18.5 (34.89%) | $9.1 \times 10^{-6}$ (37.1%) | 166.4 (37.1%) | n = 10 |
| | <sup>56</sup>Fe | 39.6 (31.51%) | $2.0 \times 10^{-5}$ (33.5%) | 350.3 (33.5%) | n = 10 |
| | <sup>63</sup>Cu | 0.9 (32.783%) | $4.7 \times 10^{-7}$ (35.6%) | 7.4 (35.6%) | n = 10 |
| | <sup>66</sup>Zn | 39.1 (32.69%) | $1.9 \times 10^{-5}$ (31.7%) | 292.8 (31.7%) | n = 10 |
| Petrolatum-Embedded | <sup>31</sup>P | 17044.8 (37.50%) | $9.4 \times 10^{-3}$ (39.8%) | $3.0 \times 10^5$ (39.8%) | n = 8<sup>b</sup> |
| | <sup>55</sup>Mn | 29.3 (24.48%) | $1.6 \times 10^{-5}$ (36.5%) | 298.5 (36.5%) | n = 8 |
| | <sup>56</sup>Fe | 38.2 (20.67%) | $2.1 \times 10^{-5}$ (24.1%) | 376.5 (24.1%) | n = 8 |
| | <sup>63</sup>Cu | 1.6 (45.31%) | $8.6 \times 10^{-7}$ (41.3%) | 13.7 (41.3%) | n = 8 |
| | <sup>66</sup>Zn | 50.2 (18.42%) | $2.8 \times 10^{-5}$ (24.3%) | 421.2 (24.3%) | n = 8 |
| Petrolatum-Coated | <sup>31</sup>P | 80289.6 (78.34%)<br>20738.3 (15.21%) | $3.8 \times 10^{-2}$ (90.0%)<sup>c</sup><br>$6.6 \times 10^{-3}$ (12.7%)<sup>d</sup> | $1.2 \times 10^6$ (90.0%)<sup>c</sup><br>$2.1 \times 10^5$ (12.7%)<sup>d</sup> | n = 10<br>n = 4 |
| | <sup>55</sup>Mn | 43.6 (48.40%) | $1.7 \times 10^{-5}$ (30.6%) | 304.8 (30.6%) | n = 10 |
| | <sup>56</sup>Fe | 53.8 (15.63%) | $2.2 \times 10^{-5}$ (21.7%) | 396.7 (21.7%) | n = 10 |
| | <sup>63</sup>Cu | 2.9 (35.35%) | $1.2 \times 10^{-6}$ (39.8%) | 19.1 (39.8%) | n = 10 |
| | <sup>66</sup>Zn | 71.8 (30.14%) | $2.9 \times 10^{-5}$ (32.4%) | 446.3 (32.4%) | n = 10 |

<sup>a</sup>Measured metal content in LA-ICP-TOF-MS imaged worms were normalized to respective worm body volume
<sup>b</sup>Two worms were not completely ablated and omitted from total element content calculations
<sup>c</sup>Phosphorus content altered due to residual M9 buffer on imaged worms from humid working conditions
<sup>d</sup>Subset of worms that did not have residual M9 due to high humidity

MANOVA analysis of metal profiles from *C. elegans* prepared from different embedding media workflows showed statistically significant differences in total metal content between preparation techniques **(Figure 4A-B)**. Post hoc linear discriminant analysis (LDA) revealed that both petrolatum-based methods clustered similarly, indicating that these two groups were not statistically different from each other. Interestingly, a multivariate outlier appeared in the petrolatum-coated *C. elegans* group that did not appear as an outlier from preliminary univariate investigations **(Figure 4B)**. Evaluation of the LDA model determined the macro-AUC of the training and test splits as 0.957, and 0.963, suggesting that the model performed well, and that the observed outlier is a product of extreme biological variance **(Figure S17)**. OCT-embedded *C. elegans* were found clustered separately from the petrolatum methods, confirming that OCT-embedded *C. elegans* were the statistically significant group identified from MANOVA **(Figure 4B)**. When quantified using gelatin standards, OCT-embedded *C. elegans* yielded the least signal for all elements when compared to both petrolatum-based methods (Wilks’ Λ = 0.2659, *F* (8, 44) = 5.1651, p < 0.001), indicating that the residual film on dried *C. elegans* may suppress plume formation during laser ablation. While no statistical significance between petrolatum-based methods was observed, previous imaging results showed that petrolatum-embedded *C. elegans* suffered from inconsistent ablation whereas petrolatum-coated *C. elegans* did not experience this. Additionally, imaging quality appeared different between these two sample preparation techniques. A larger sample set is necessary to confirm the observed differences between petrolatum-embedded and petrolatum-coated imaging. Furthermore, the accuracy of the quantitative imaging data must be confirmed against validated synchrotron-XFM imaging or ICP-QQQ-MS data with worms of the same developmental stage.

**Figure 4:**
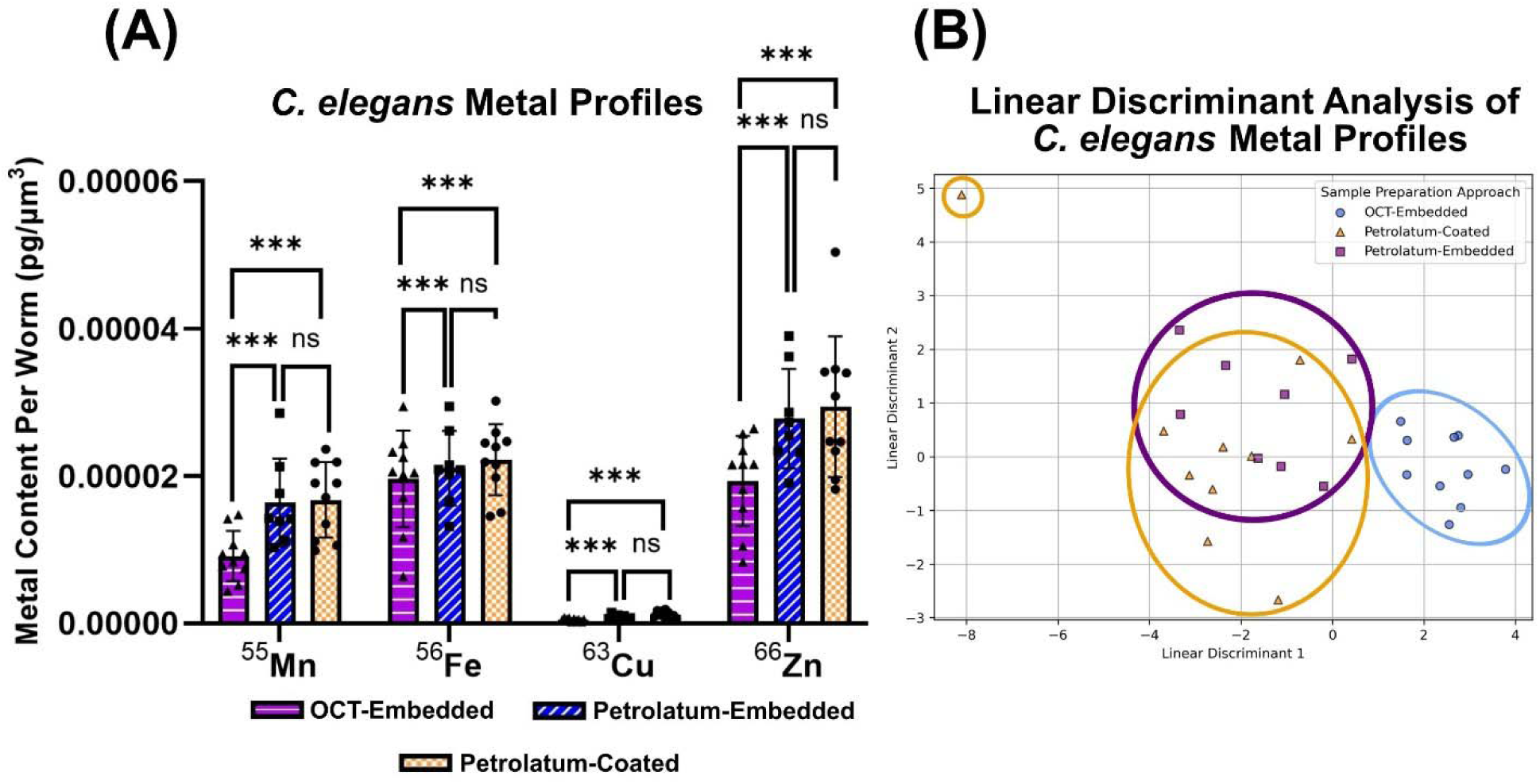
MANOVA analysis of total metal content in *C. elegans* from quantitative LA-ICP-TOF-MS imaging prepared using different embedding techniques. (A) Volume-normalized metal content from *C. elegans* visualizing statistically significant differences in metal content between sample preparation techniques from MANOVA and post hoc testing. Different colored and patterned bars represent the total metal content of worms for ^55^Mn, ^56^Fe, ^63^Cu, and ^66^Zn. Solid shapes indicate individual replicates (*n* = 10 for OCT-embedded, *n* =8 for petrolatum-embedded, *n =*10 for petrolatum-coated worms). One-way MANOVA (Wilks’ Λ = 0.2431, *F* (8, 44) = 5.655, p < 0.001). (B) Linear Discriminant Analysis of z-score normalized *C. elegans* metal profiles visualizing both petrolatum-based methods cluster similarly while OCT-embedded *C. elegans* are significantly different and cluster separately. OCT-embedded data points are represented by blue circles, petrolatum-embedded data points are represented by purple squares, and petrolatum-coated are represented by orange triangles. ***= p<0.001.

## Conclusion

Each sample preparation approach for LA-ICP-TOF-MS imaging of *C. elegans* offers different advantages and limitations and is summarized graphically in **Figure 5**. OCT-embedded and petrolatum are commercially available and elementally pure for the purpose of quantitative elemental imaging applications. OCT-embedded offers the highest quality microscopy imaging with minimal non-specific fluorescence, however, as a water based embedding media it desiccates rapidly which impact *C. elegans* morphology and metal localization. Petrolatum coating offers consistent ablation with moderate sample preservation qualities allowing for preservation up to 8 hours at ambient conditions. Non-specific fluorescence remains an issue with the petrolatum coating method which prevents multi-modal imaging applications using fluorescence microscopy. Petrolatum embedding offers the longest sample preservation of the explored workflows. The non-specific fluorescence in petrolatum-embedded worms can be filtered out through adjusting microscopy parameters, while in petrolatum-coated worms the potentially scattered fluorescence cannot be corrected.

**Figure 5:**
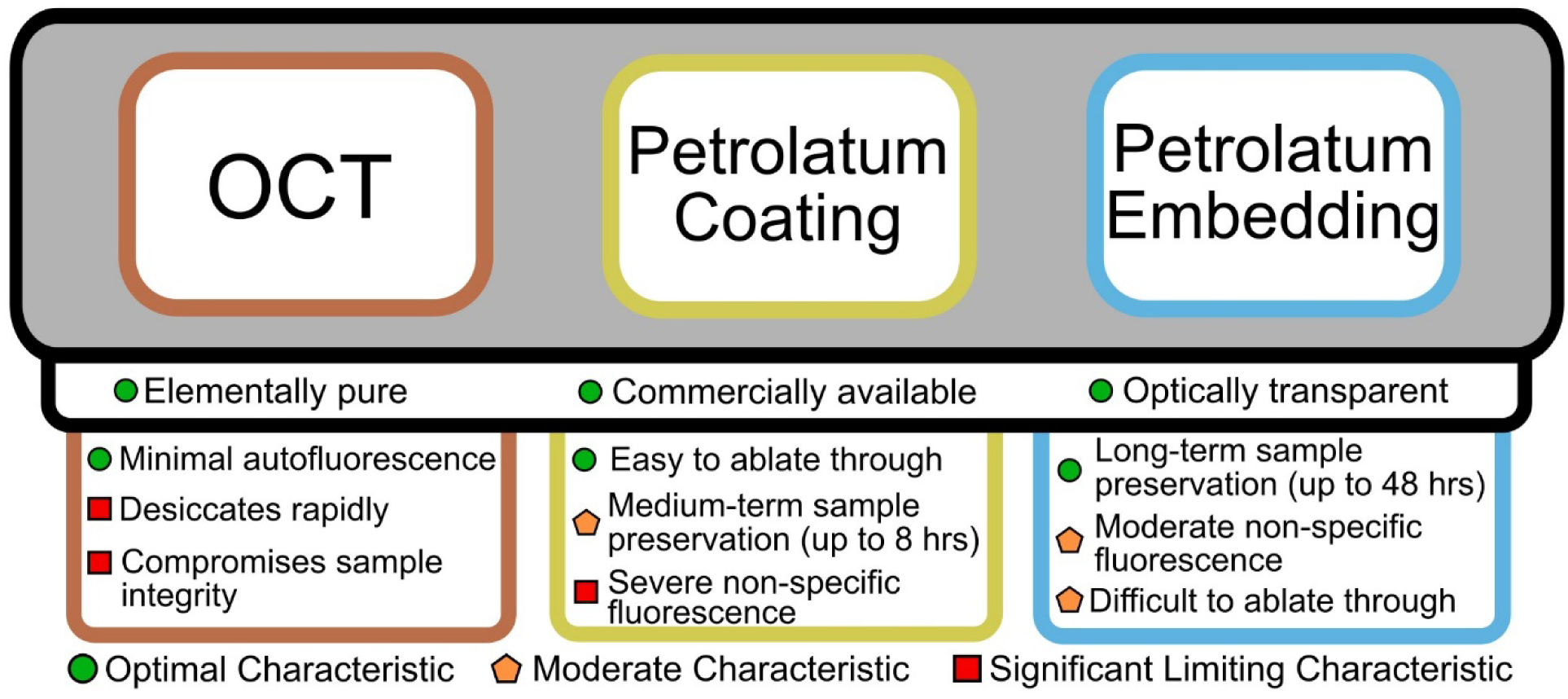
Summary of different tested embedding approaches for microscopy and complimentary LA-ICP-TOF-MS imaging of *C. elegans*. Shared characteristics between different preparation techniques are found in black bordered boxes that overlap multiple technique names. Characteristics unique to a specific preparation technique are listed below in the colored bordered boxes. Green circles indicate the characteristic is highly desirable, orange pentagons indicate that the characteristic is not inherently optimal but manageable and acceptable, and red squares indicate that the characteristic is a significant limiting factor that cannot be overcome to optimize the preparation technique.

While quantitation was performed using gelatin standards, these standards were not matrix-matched to reflect the true ablation characteristics of embedded *C. elegans*. Additional steps in modifying the properties of gelatin standards can be performed to better match the sample matrix, however, this requires complimentary ICP-QQQ-MS or synchrotron-XFM quantitation to confirm the true concentrations of imaged samples.^13^ With these complimentary techniques, future research will require optimizing gelatin standard properties, gelatin thickness, and calibration curve concentration ranges.

LA-ICP-MS imaging is an important imaging modality for metal biology research in model animals. *Caenorhabditis elegans* presents a powerful animal model for fundamental biological research. Previous studies have performed LA-ICP-MS imaging on *C. elegans*, but the sample preparation and technological limitations prevented accurate and comprehensive imaging.^29–31^ Our study developed a 3D-printed tool for on-slide embedding of *C. elegans* and demonstrated that petrolatum can serve as an ideal embedding media for preserving reporter fluorescence and *C. elegans* morphology when compared to conventional embedding media such as OCT. We found that petrolatum possessed sufficient optical transparency for microscopy and was elementally pure to produce minimal background interference during LA-ICP-MS imaging.

Furthermore, we evaluated sample preservation of worms at ambient conditions when prepared by OCT-embedded, petrolatum-embedded, and petrolatum-coated which showed each approach yielded different qualities of brightfield and fluorescence imaging. Through over-sampling laser shots we were able to achieve an effective 2-μm spatial resolution, allowing for identification of cellular elemental compartmentalization that has been previously only observed in XFM studies.^48,61^ We further established a data analysis pipeline on this over-sampling method towards quantitative LA-ICP-TOF-MS imaging using gelatin standards for single-whole worm elemental quantitation. Using LA-ICP coupled with a TOF analyzer, we successfully demonstrated quantitative imaging of most endogenous elements simultaneously in one sample, allowing co-localization of worm fluorescence reporters, brightfield images and different elements. In summary, we developed a comprehensive sample preparation, instrumental analysis and data processing workflow to enable high-resolution quantitative LA-ICP-TOF-MS imaging workflow on *C. elegans*, providing a powerful tool for future metal biology research.

## Author Contributions

A.J.R, A.S –conceptualization; A.J.R. –experimental design, 3D-printed tool design, embedding method development and validation, microscopy, quantitative LA-ICP-TOF-MS imaging experiments, ICP-QQQ-MS experiments, data analysis pipeline, statistics, data visualization, writing – original draft; A.S –experimental design, preliminary embedding method validation, ICP-QQQ-MS experiments, data interpretation, writing – review and editing; K.M – ICP-QQQ-MS experiments, experimental design, writing – review and editing; T.V.O – conceptualization, experimental design, writing – review and editing, funding acquisition, supervision; T.A.Q – conceptualization, experimental design, data interpretation, writing – review and editing, funding acquisition, supervision.

## Notes

The authors declare that there are no conflicts of interest.

## Data Availability Statement

The data that support the findings of this study are available in the Methods, Results, and Supplemental Material of this article. Other original datasets are available upon request to the corresponding authors.

## Acknowledgements

The authors gratefully acknowledge support from Michigan State University start-up funds to T.A.Q. and NIH grants R01GM115848 and R01GM038784 to T.V.O. This research was carried out in collaboration with the Quantitative Bio Element Analysis and Mapping (QBEAM) Center at Michigan State University and The National Research Resource for Quantitative Elemental Mapping for the Life Sciences (QE-Map) under Grants P41 GM135018 and S10OD026786 to T.V.O. from the National Institutes of Health. All *C. elegans* and OP50 strains used in this work were provided by the CGC, which is funded by NIH Office of Research Infrastructure Programs (P40 OD010440). We thank Daniel Scsavnicki and Ryutaro Einar Jacobson for providing 3D-printing equipment to produce initial drafts of the tools used in this research. We thank Santosh Patnaik, MD, PhD for creating the ImageJ macro “InteredgeDistance_v1.1_ImageJMacro” and providing the macro as an open-source tool for ImageJ users.

