## Supplemental for "On-slide Preparation of *Caenorhabditis elegans* Towards Quantitative, High-Resolution LA-ICP-TOF Mass Spectrometry Imaging"

### Table of Contents

**Table S1:** Anycubic Photon Mono 4 3D printing parameters
**Table S2:** Microscopy imaging parameters
**Table S3:** LA-ICP-TOF imaging parameters
**Table S4:** Measured Gelatin Standard Concentrations
**Table S5:** Calculated limits of detection and quantitation
**Figure S1**: *C. elegans* dimension analysis and workflow for determining appropriate gelatin standard thickness
**Figure S2:** Microscopy imaging of *C. elegans* embedded in alternative media
**Figure S3:** Petrolatum preparation and properties under microscopy observations
**Figure S4:** Petrolatum-embedded microscopy time-series imaging
**Figure S5:** Anatomical feature assignment to petrolatum-embedded *C. elegans*
**Figure S6:** Changes in *C. elegans* morphology due to intestinal necrosis
**Figure S7:** Petrolatum-coated microscopy time-series imaging
**Figure S8:** Elemental indicators present in microscope slides for complete sample ablation
**Figure S9:** 5 µm versus 2 µm spatial resolution LA-ICP-TOF-MS imaging
**Figure S10:** Variation in *C. elegans* embedding height in petrolatum causes inconsistent LA-ICP-TOF-MS imaging
**Figure S11:** Dimensions of *C. elegans* determined by ImageJ
**Figure S12:** Comprehensive LA-ICP-TOF-MS imaging of day 1 gravid adult Bristol N2 *C. elegans* embedded in petrolatum
**Figure S13:** Multi-day calibration curve data
**Figure S14:** Calibration Curve sensitivity drift contributes to calibration standard deviations
**Figure S15:** Evaluation of residual $^{31}$P content from M9 buffer rinsing
**Figure S16:** Unnormalized versus volume-normalized total metal content in *C. elegans*
**Figure S17:** Macro-average ROC curves evaluating LDA model performance

### Supporting Information

**Table S1:** Anycubic Photon Mono 4 3D Printer Parameters

| Slice Parameters | | | |
|---|---|---|---|
| Layers Thickness (mm) | 0.050 | Control Type | Basic |
| Normal Exposure Time (s) | 2.5 | Z Lift Distance (mm) | 6 |
| Off Time (s) | 1 | Z Lift Speed (mm/s) | 4 |
| Bottom Exposure Time (s) | 30 | Z Retract Speed (mm/s) | 6 |
| Bottom Layers | 5 | | |
| Anti-alias | 1 | | |
| Use Random Erode Shell | Off | | |

**Table S2:** Zeiss Axio Imager.Z2 Microscopy imaging parameters

| ***Brightfield parameters*** | | | ***GFP channel parameters*** | | | ***mCherry/tdTomato channel parameters*** | | |
|---|---|---|---|---|---|---|---|---|
| **Parameter** | **Unit** | **Value** | **Parameter** | **Unit** | **Value** | **Parameter** | **Unit** | **Value** |
| Lightsource intensity (TL Halogen lamp) | V | 4.2 | Lightsource intensity (Xcite Xylis lamp) | % | 15 | Lightsource intensity (Xcite Xylis lamp) | % | 15 |
| Exposure time | ms | 10.00 | Exposure time | ms | 100 | Exposure time | ms | 100 |
| Intensity | % | 30 | Intensity | % | 15 | Intensity | % | 30 |
| Black contrast | ~ | 0 | Black contrast | ~ | 200 | Black contrast | ~ | 150 |
| White contrast | ~ | 6000-9000 | White contrast | ~ | 1500 | White contrast | ~ | 750 |
| ***Camera parameters*** | | | | | | | | |
| **Parameter** | **Unit** | **Value** | | | | | | |
| Exposure time | ms | 10.00 | | | | | | |
| Auto Exposure | on/off | on | | | | | | |
| Intensity | % | 30 | | | | | | |

**Table S3:** Essential LA-ICP-TOF MS imaging parameters

| ***ICP-MS Parameters (Tofwerk S2)*** | | | ***Time-of-Flight Parameters (Tofwerk S2)*** | | | **Laser Ablation Parameters (ESL Bioimage 266 nm)** | | |
|---|---|---|---|---|---|---|---|---|
| **Parameter** | **Unit** | **Value** | **Parameter** | **Unit** | **Value** | **Parameter** | **Unit** | **Value** |
| **RF Power** | W | 1550 | **m/z range** | amu | 14 - 256 | **Spot Size** | µm | 5 |
| **Sampling Depth** | mm | 4.9 | **Resolution** | m/Δm | 1000 | **Interline Distance (y)** | µm | 2 |
| **Cone Material** | Nickel | - | **ODG Settings** | ms | 20 | **Overlap (x)** | µm | 3 |
| **Cone Insert (STD)** | mm | 3.5 | **Notch Bias** | V | -80 | **Repetition Rate** | Hz | 125-200 |
| **Plasma Gas Flow** | L/min | 14.0 | **Notch (40 amu)** | V | 3 | **Laser Power** | % | 70 |
| **Auxillary Gas Flow** | L/min | 8.0 | **Notch (36 amu)** | V | 1 | **Laser Fluence** | J/cm$^2$ | 9.5-10.2 |
| **Nebulizer Gas Flow** | L/min | 0.9 - 1.0 | **Notch (28 amu)** | V | 2 | **Sample Energy** | mJ | .005 |
| **Measurement Mode** | CCT Mode | - | **Notch (15.8 amu)** | V | 3 | **Imaging Cup Flow Rate (He)** | mL/min | 200 |
| **CCT Gas Flow (100% He)** | mL/min | 5 | | | | **Imaging Chamber Flow Rate (He)** | mL/min | 250 |
| **CCT Focus lens** | V | -19.5 | | | | **PEEK Tubing I.D.** | mm | 0.75 |
| **CCT Entry Lens** | V | -180 | | | | | | |
| **CCT Mass** | V | 250 | | | | | | |
| **CCT Bias** | V | -9 | | | | | | |
| **CCT Exit Lens** | V | -200 | | | | | | |

### Supporting Information

**Table S4:** Essential element concentrations in gelatin standards measured by ICP-QQQ-MS

| Standard | $^{31}$P | $^{55}$Mn | $^{56}$Fe | $^{63}$Cu | $^{66}$Zn |
|---|---|---|---|---|---|
| Gelatin 50 ppm | ~ | 50.18 | 51.50 | 50.59 | 50.45 |
| Gelatin 25 ppm | ~ | 24.06 | 24.30 | 23.49 | 23.47 |
| Gelatin 12.5 ppm | ~ | 11.37 | 11.73 | 11.18 | 11.61 |
| Gelatin 4000 ppm | 3798.42 | ~ | ~ | ~ | ~ |
| Gelatin 2000 ppm | 1870.44 | ~ | ~ | ~ | ~ |
| Gelatin 1000 ppm | 945.54 | ~ | ~ | ~ | ~ |
| Gelatin Blank | 3.29 | 0.02 | 0.56 | 0.026 | 0.74 |

**Table S5: Quantitative LA-ICP-TOF-MS Limits of Detection and Quantitation**

**Calculated LODs and LOQs reported as parts per million (ppm)**

| Imaging Experiment | | | Element $^{31}$P | $^{55}$Mn | $^{56}$Fe | $^{63}$Cu | $^{66}$Zn |
|---|---|---|---|---|---|---|---|
| OCT-Embedded | 1 | LOD | 11.44 | 0.04 | 0.41 | 0.04 | 0.13 |
| | | LOQ | 34.32 | 0.12 | 1.22 | 0.13 | 0.38 |
| OCT-Embedded | 2 | LOD | 20.03 | 0.03 | 0.19 | 0.03 | 0.07 |
| | | LOQ | 60.10 | 0.10 | 0.57 | 0.08 | 0.21 |
| Petrolatum-Embedded | 3 | LOD | 17.19 | 0.10 | 0.47 | 0.08 | 0.33 |
| | | LOQ | 51.57 | 0.29 | 1.42 | 0.24 | 0.99 |
| Petrolatum-Embedded | 4 | LOD | 49.47 | 0.06 | 1.42 | 0.11 | 3.38 |
| | | LOQ | 148.41 | 0.19 | 4.26 | 0.34 | 10.14 |
| Petrolatum-Embedded | 5 | LOD | 118.11 | 0.09 | 0.36 | 0.07 | 0.29 |
| | | LOQ | 354.32 | 0.26 | 1.08 | 0.20 | 0.87 |
| Petrolatum-Coated | 6 | LOD | 42.06 | 0.06 | 0.56 | 0.07 | 0.16 |
| | | LOQ | 126.18 | 0.19 | 1.67 | 0.20 | 0.48 |
| Petrolatum-Coated | 7 | LOD | 426.04 | 0.11 | 1.33 | 0.25 | 0.81 |
| | | LOQ | 1278.12 | 0.34 | 4.00 | 0.74 | 2.42 |
| Petrolatum-Coated | 8 | LOD | 9.46 | 0.06 | 0.39 | 0.05 | 0.17 |
| | | LOQ | 28.37 | 0.17 | 1.17 | 0.14 | 0.51 |
| Averages (RSD%) | | LOD | 86.72 (163.3%) | 0.07 (40.1%) | 0.64 (72.7%) | 0.09 (81.8%) | 0.67 (167.8%) |
| | | LOQ | 260.17 (163.3%) | 0.21 (40.1%) | 1.92 (72.7%) | 0.26 (81.8%) | 2.00 (167.8%) |

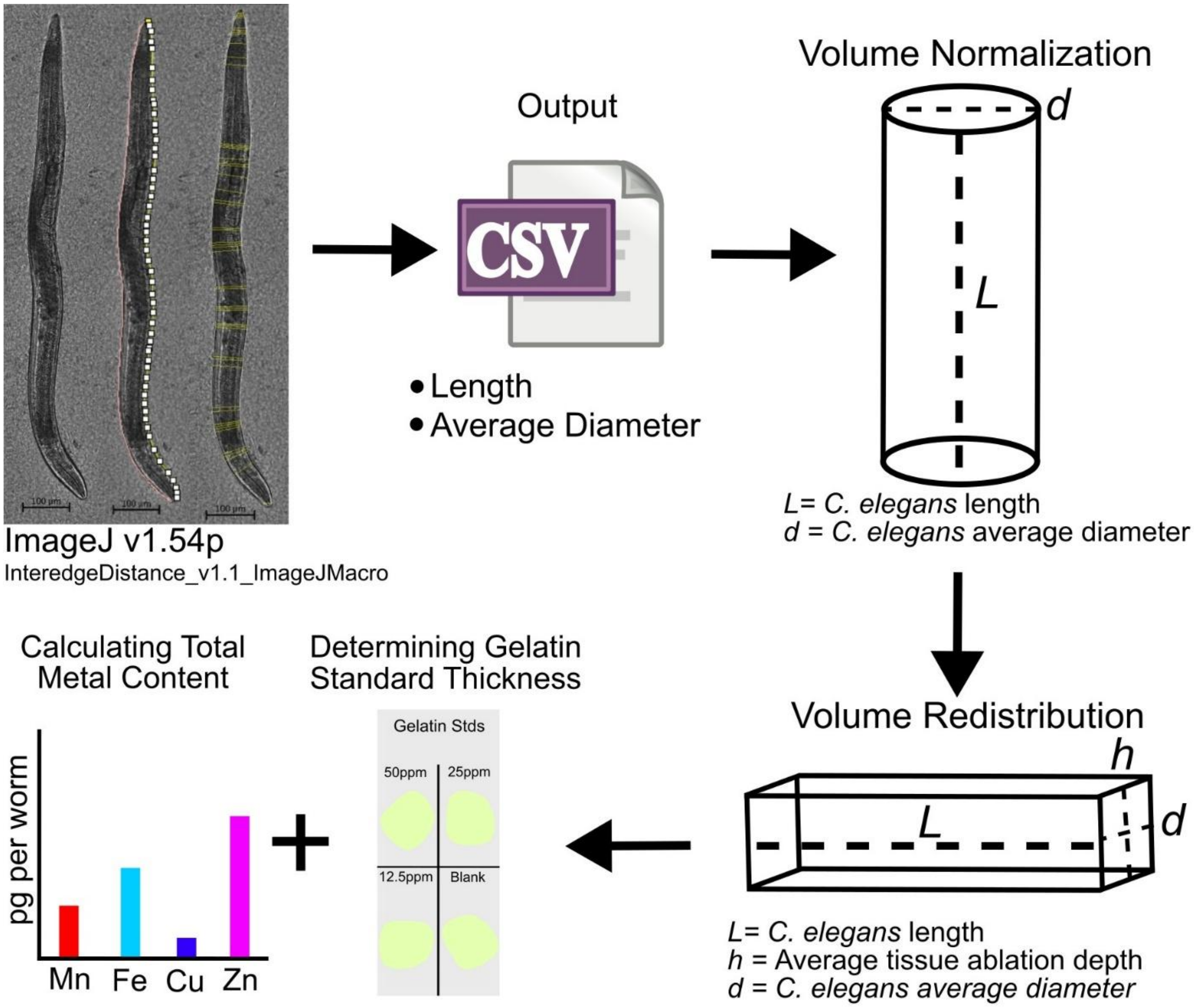


**Figure S1:** Workflow diagram for determining the dimensions of imaged *C. elegans* using ImageJ and the macro InteredgeDistance_v1.1_ImageJMacro. Microscopy images of worms are calibrated and imported into ImageJ where the user defines the outline of the sample along the left and right sides of the worm. The length of the worm is measured, and the diameter of the worm is sampled at up to 50 different positions along the user defined lines, generating an average diameter. *C. elegans* volume is determined from the measured lengths and diameters and the volume of the worm is normalized to the shape of cylinder. The volume is then redistributed from a cylinder to a cuboid to determine the average thickness of the worm. The average thickness of the worm is then used for downstream processes including gelatin standard thickness determination and total metal content extraction from LA-ICP-TOF-MS images.

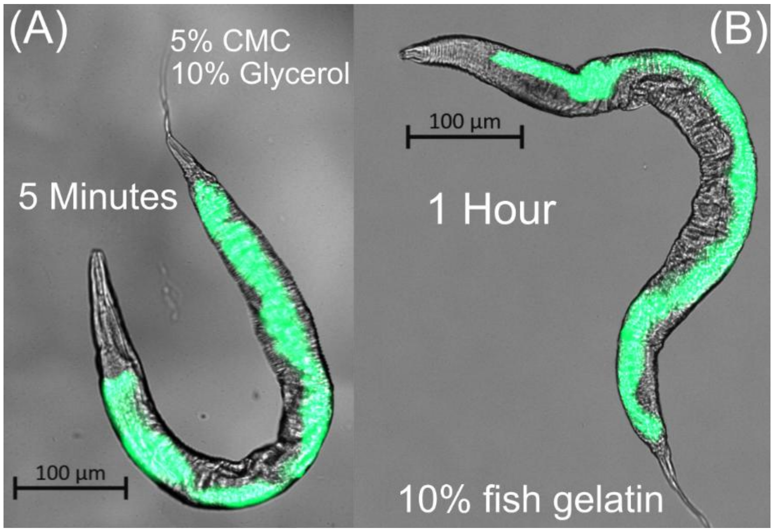


**Figure S2:** Brightfield and GFP imaging of a gravid adult day 1 SJ4144 *C.* elegans embedded in alternative water-based embedding media. (A) A day 1 gravid SJ4144 *C.* elegans in 5% carboxymethyl cellulose (CMC) spiked with 10% glycerol. (B) A day 1 gravid SJ4144 *C.* elegans in 10% fish gelatin. GFP expression visualizes the intestine of *C. elegans*. The wrinkled cuticle in the brightfield channel indicates desiccation. Times indicate the earliest point when complete desiccation occurred. Black scale bars indicate 100 µm.

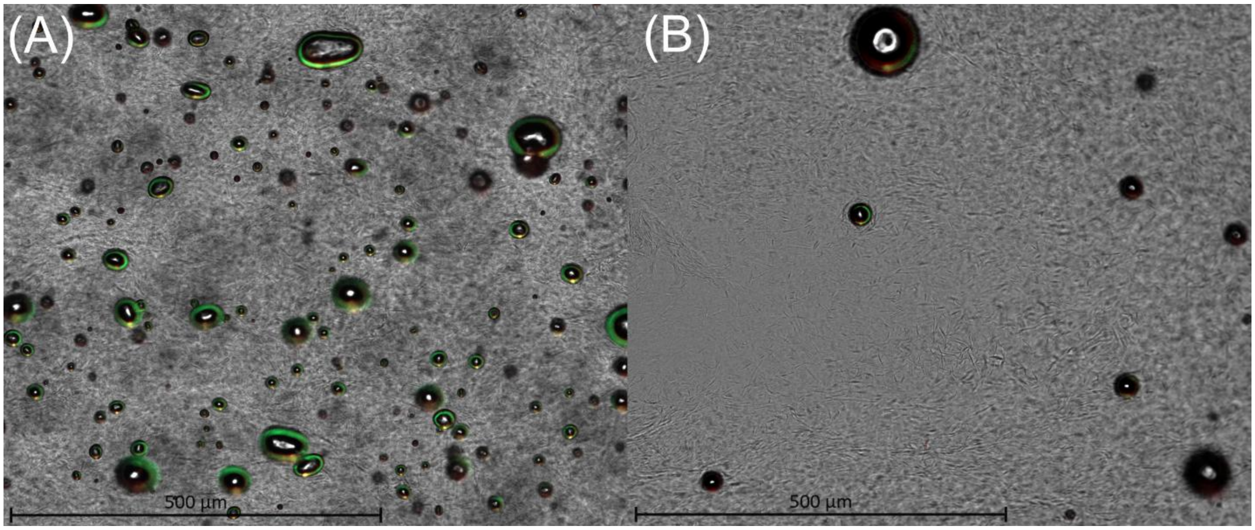


**Figure S3:** Petrolatum characteristics viewed under microscope. (A) Initial petrolatum application directly from Vaseline container, (B) and pre-heated and degassed petrolatum application showing that heating and sonicating petrolatum prior to application can reduce bubble formation during the embedded process. Black scale bars indicate 500 µm.

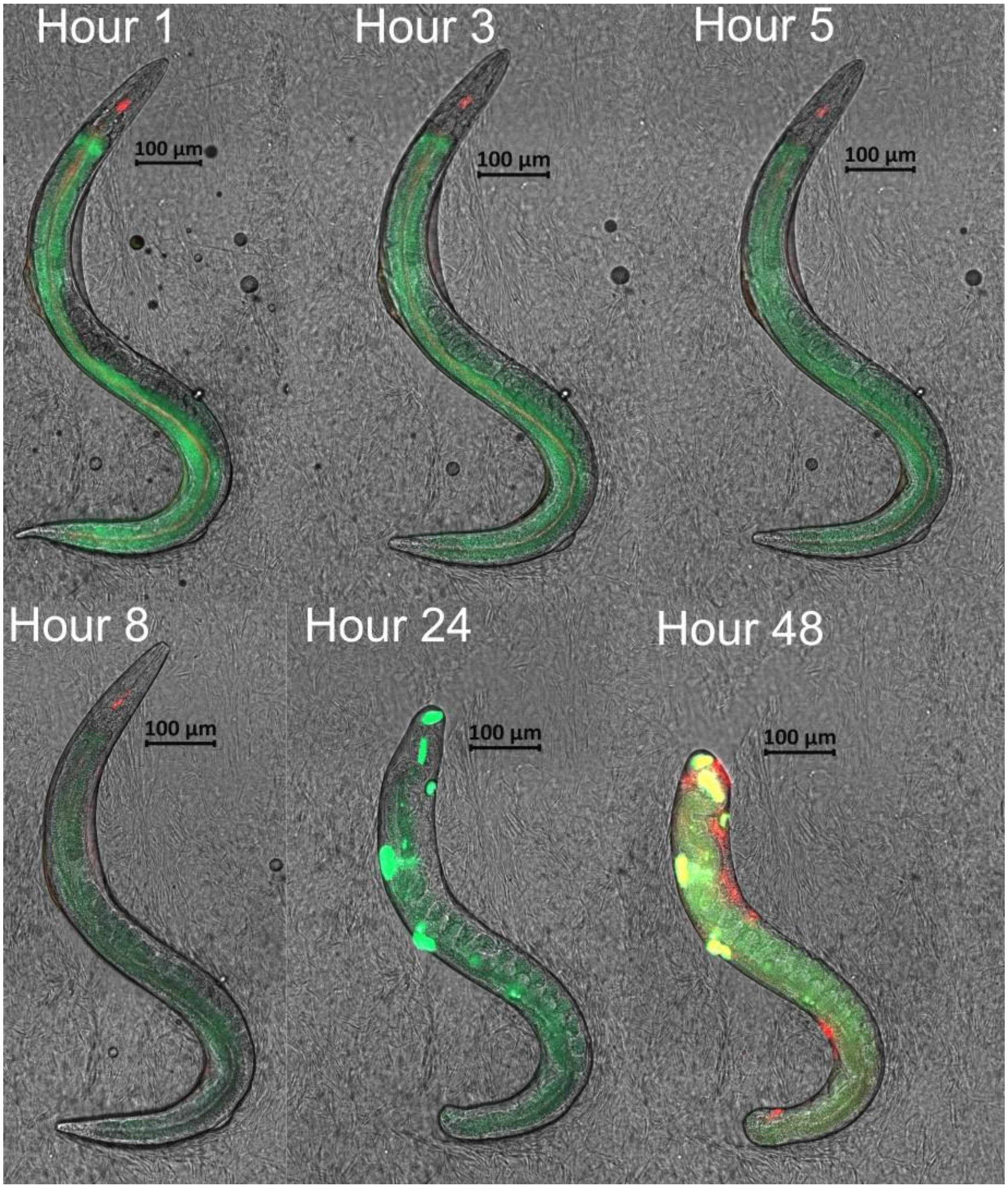


**Figure S4:** Evaluation of petrolatum for day 1 gravid adult SJ4144 *C. elegans* preservation and 48-hr tracking on post-processing preservation at ambient conditions. Petrolatum-embedded worms were sufficiently preserved such that tissue decay occurred prior to desiccation. Green channel fluorescence imaging reveals intestinally expressed cytosolic GFP. Red channel fluorescence imaging reveals the localization of tdTomato from bacterial matter within *C. elegans*. Black scale bars indicate 100 µm.

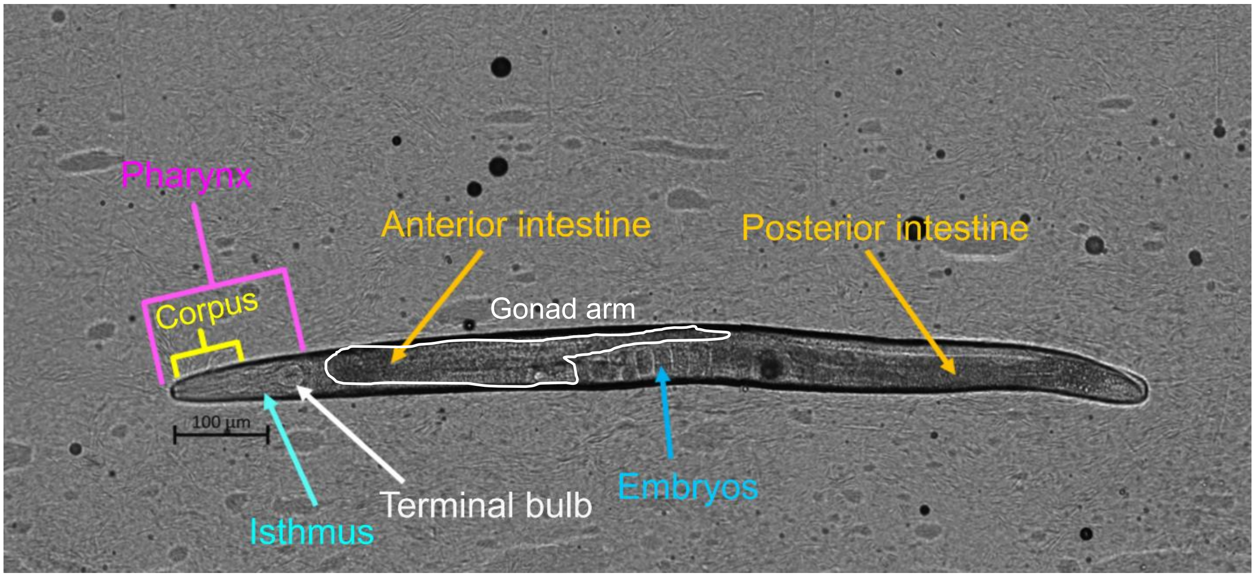


**Figure S5:** Identification of major anatomical features in an adult hermaphroditic *C. elegans* embedded in petrolatum. The pharynx is identified by the pink brackets with pharyngeal components labeled with the white, cyan and yellow indicators. The intestinal tract is identified by the orange indicators and embryos are identified in the blue indicator. Black scale bar indicate 100 µm.

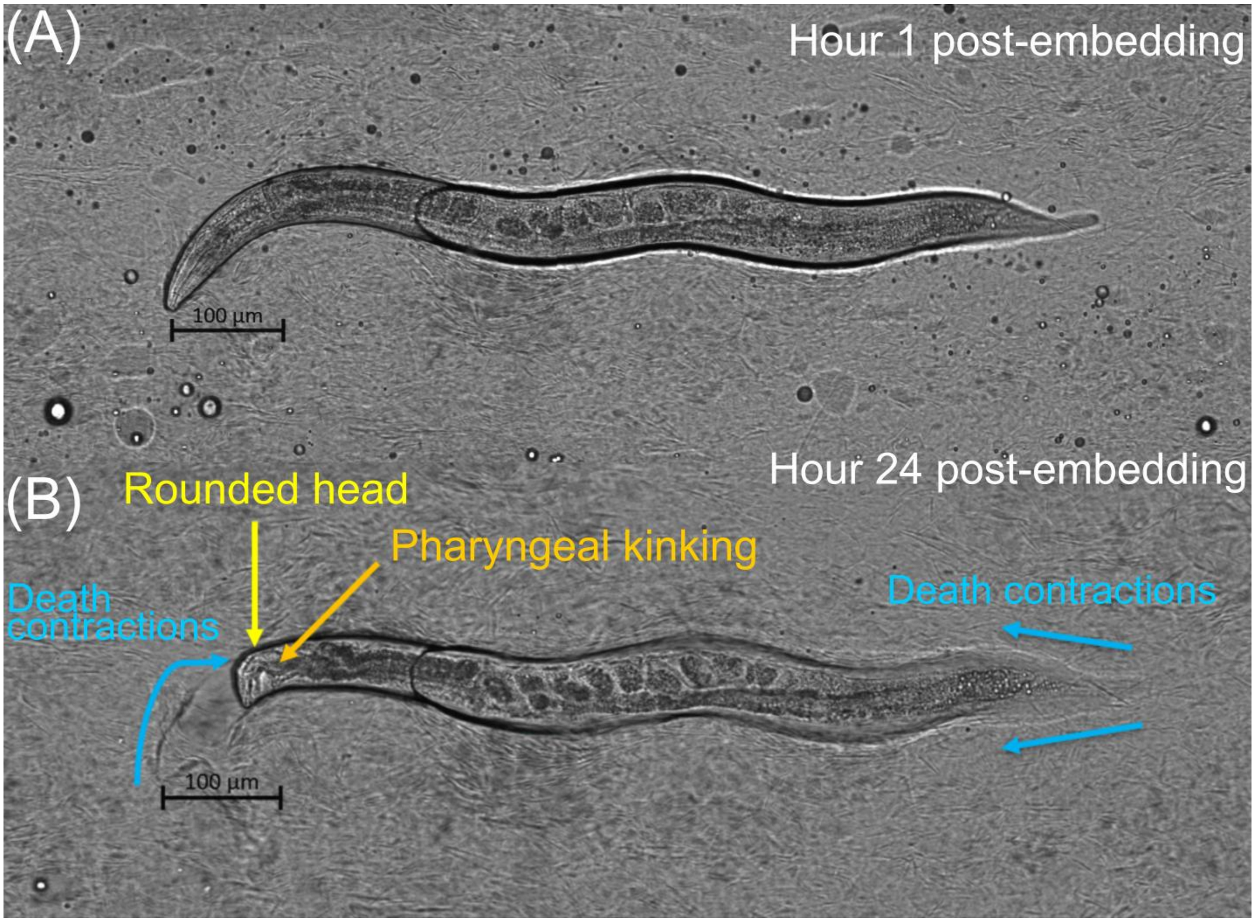


**Figure S6:** Investigating sample integrity and morphology changes of *C. elegans* embedded in petrolatum. (A) Initial embedding imaging of gravid day 1 adult *C. elegans* at the 1-hour post-embedding time point reveals no disturbed morphology of the worm. (B) Death contractions and pharyngeal kinking indicative of intestinal necrosis observed at the 24-hour post-embedding time point. Commonly observed morphological changes are indicated in blue, yellow and orange. Black scale bars indicate 100 µm.

### Supporting Information

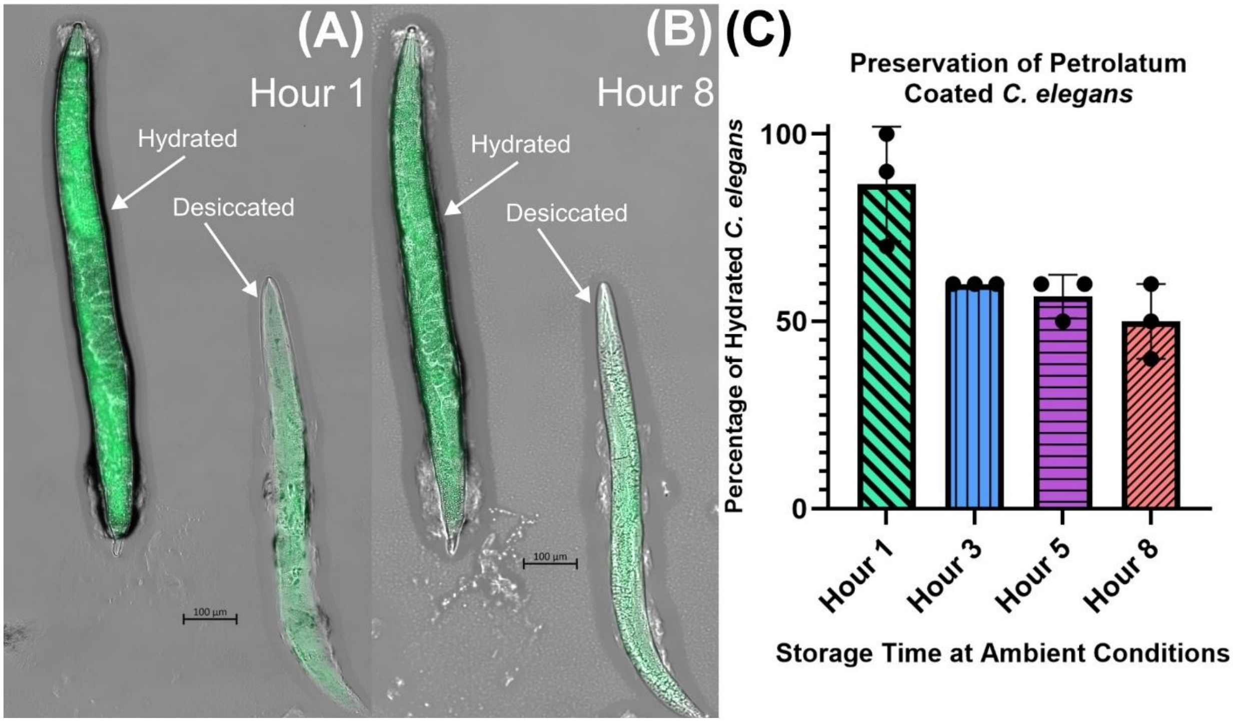


**Figure S7:** Time-series evaluation of preservation capabilities of petrolatum-coated *C. elegans*. (A) Microscopy imaging of two gravid day 1 adult SJ4144 *C. elegans* at the 1-hour post-coating time point, (B) and at the 8-hour post-coating time point. Hydrated and desiccated worms are indicated by white arrows. Immediate desiccation of coated worms is due to unintentional removal of petrolatum coating during slide transfer. (C) *C. elegans* preservation efficacy over an 8-hour period at ambient conditions. Different colored and patterned bars represent the number of hydrated worms normalized to a percentage value. Solid black circles indicate the number of experimental replicates (n = 3). Each experimental replicate imaged 10 individual worms for individual replicates n = 30 across all experiment trials Black scale bars indicate 100 µm.

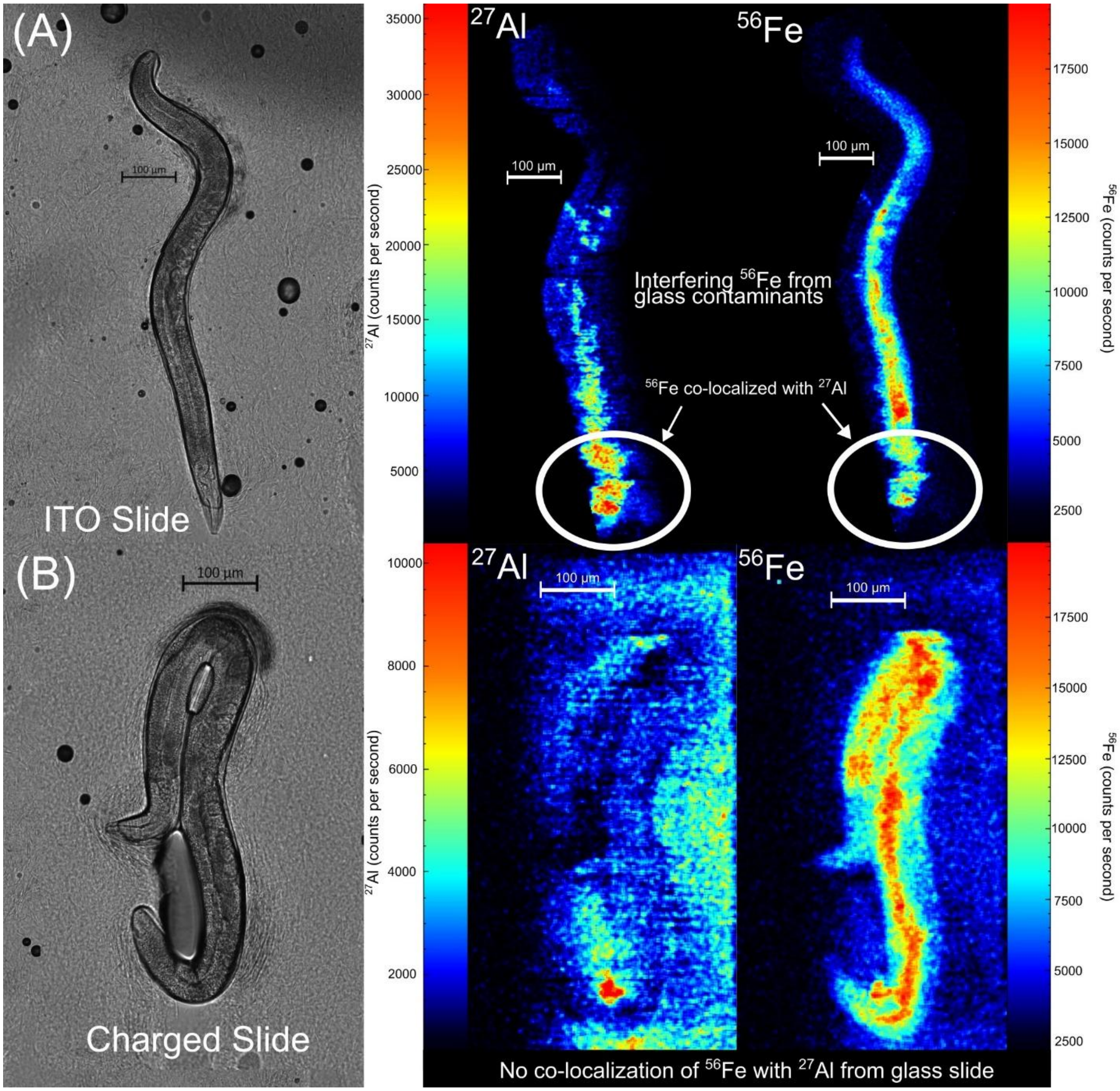


**Figure S8:** Determining optimal microscope slide for LA-ICP-TOF-MS imaging of *C. elegans*, and establishing elemental signals indicative of complete ablation of *C. elegans*. (A) Microscopy and LA-ICP-TOF imaging of a day 1 gravid adult wildtype *C. elegans* visualizing $^{27}Al$ and $^{56}Fe$ on an ITO slide producing excessive interfering $^{56}Fe$ signals indicated by the white circles co-localized to $^{27}Al$ present in the ITO slide. (B)Microscopy and LA-ICP-TOF imaging of a day 1 gravid adult wildtype *C. elegans* visualizing $^{27}Al$ and $^{56}Fe$ using an Epdredia charged slide which does not introduce excessive interfering signals from glass slide contaminants. Black scale bars for microscopy images and white scale bars in ion images indicate 100 µm.

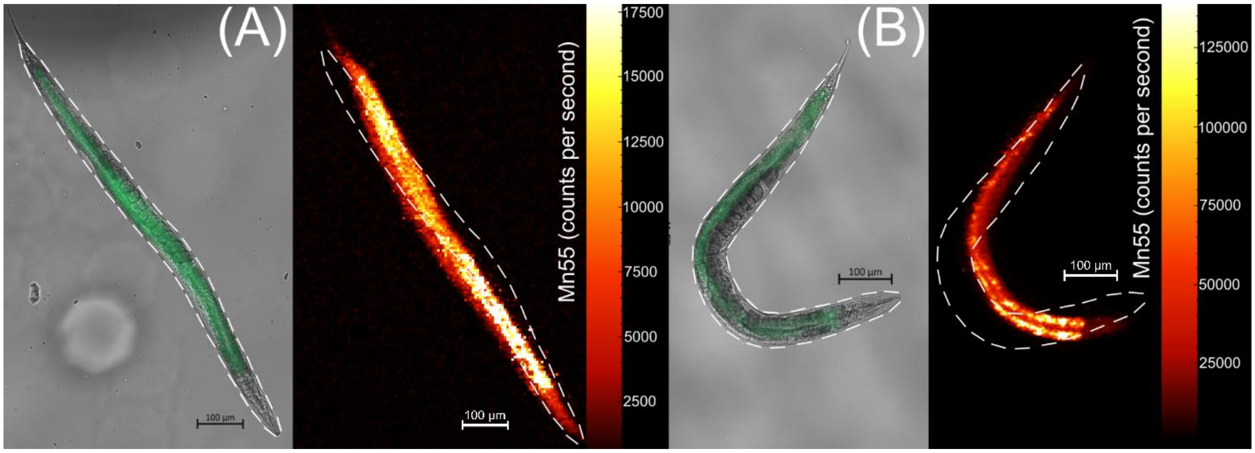


**Figure S9:** Comparison of laser spot sizes and spatial resolution quality from imaged *C. elegans*. (A) LA-ICP-TOF-MS imaging of day 1 gravid adult SJ4144 *C. elegans* using a 5 µm spot size laser profile, (B) and an effective 2 µm spot size achieved by over-sampling a 5 µm spot size laser profile showing $^{55}$Mn distribution. The effective 2 µm spot size achieves a spatial resolution capable of visualizing the intestinal lumen and differentiating intestinal cells. Outlines of imaged *C. elegans* is denoted by the white dashed lines which show that *C. elegans* embedded in OCT change shape and experience morphological changes during LA-ICP-TOF-MS imaging. Black scale bars for microscopy images and white scale bars in ion images indicate 100 µm.

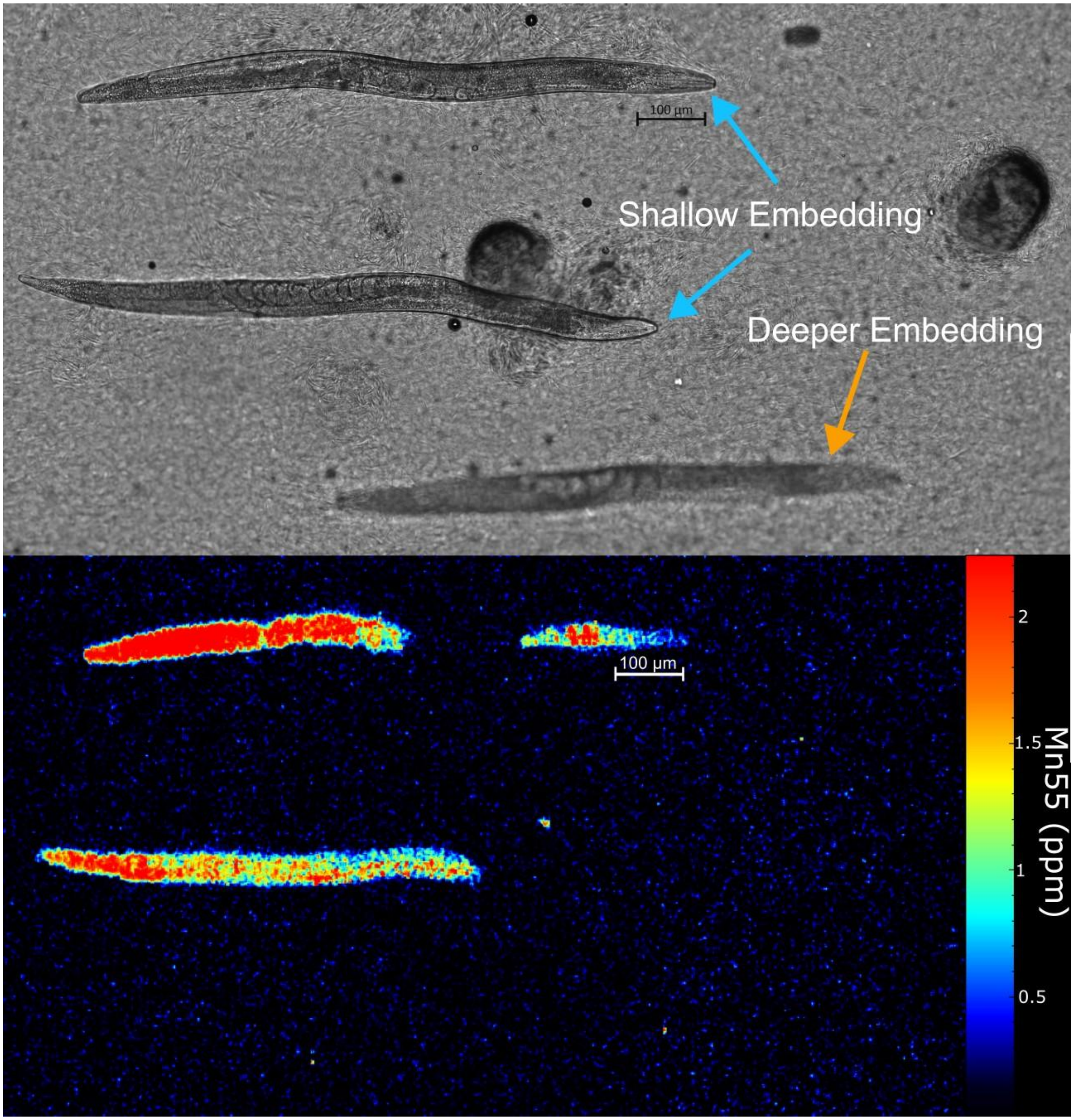


**Figure S10:** Vaseline shows laser ablation inconsistencies during LA-ICP-TOF-MS imaging. Producing in a ~100 µm embedding layer induces too much variation in *C. elegans'* vertical positioning, requiring further layering height optimization to improve imaging consistency of petrolatum-embedded worms. Black scale bars for microscopy images and white scale bars in ion images indicate 100 µm.

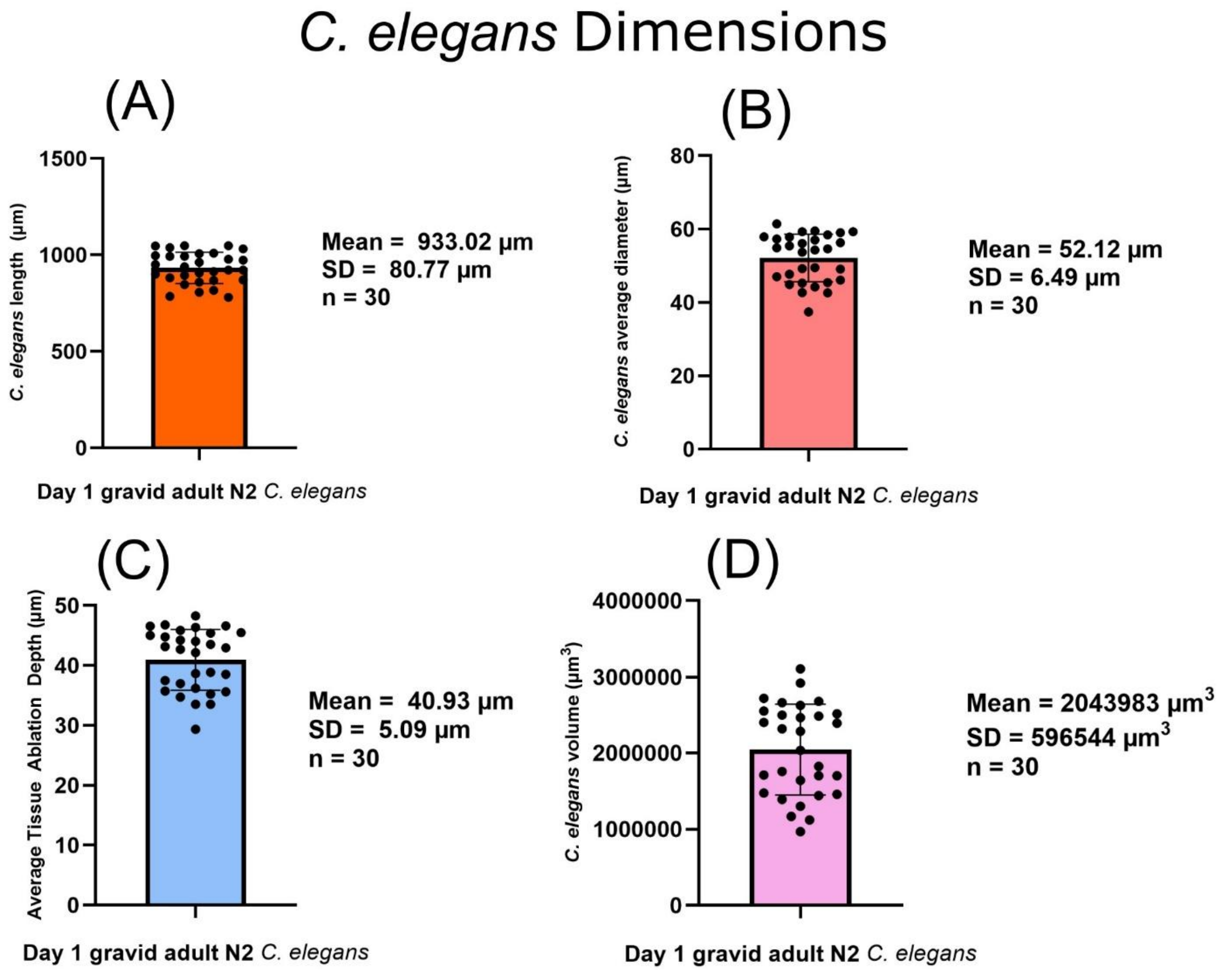


**Figure S11:** Dimensional analysis of *C. elegans* using the InteredgeDistance macro for ImageJ. (A) Day 1 adult N2 *C. elegans* were sampled within 4 hours of entering adulthood with their length, (B) average diameter across the whole body, (C) average tissue ablation depth, (D) and volumes measured by ImageJ. Average tissue ablation depth was used as reference for determining the gelatin standard sectioning thickness for quantitative LA-ICP-TOF-MS imaging. Dimensions were acquired from $n$ = 30 *C. elegans* embedded in petrolatum.

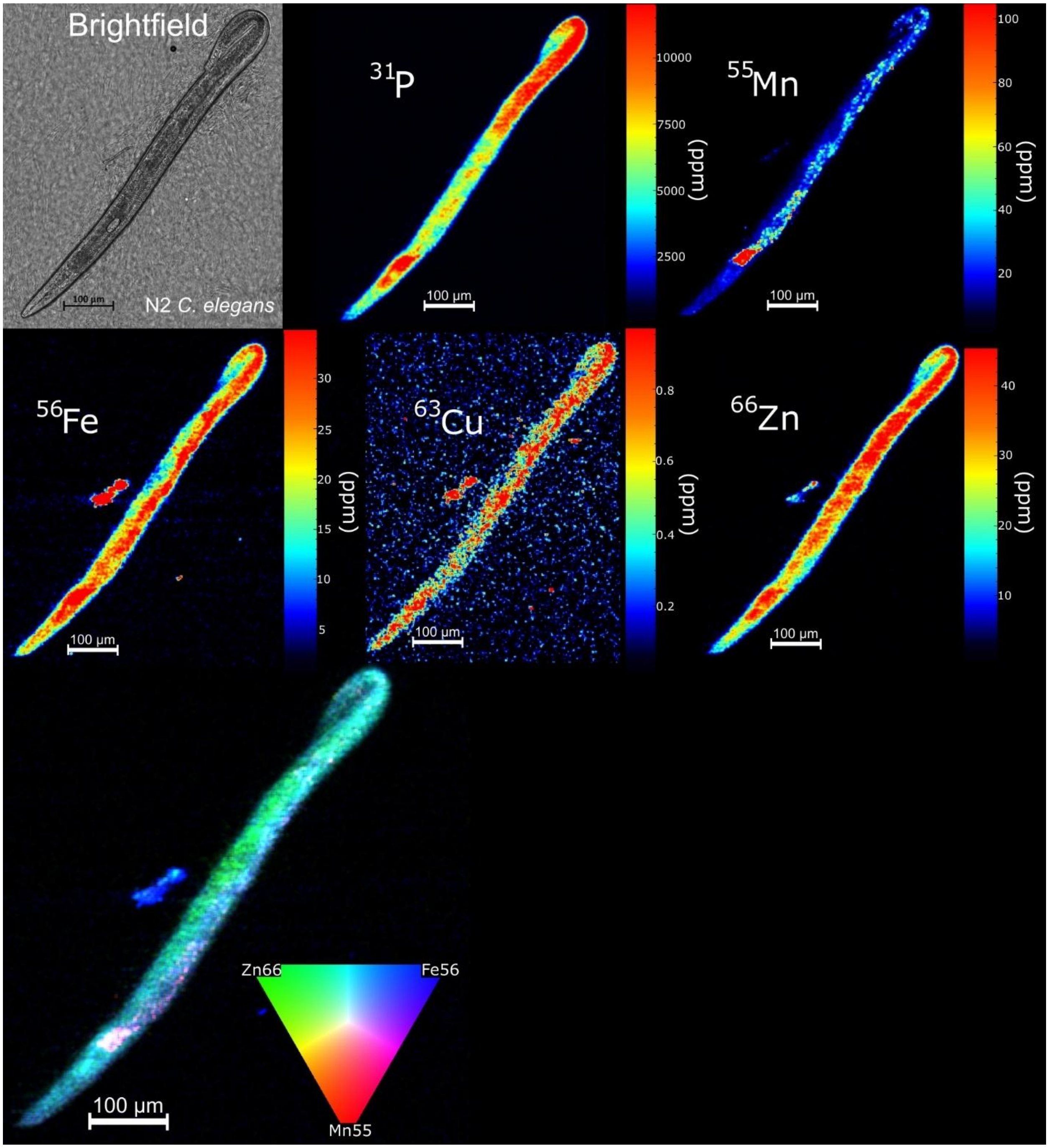


**Figure S12:** Brightfield imaging of a petrolatum-embedded day 1 gravid adult wildtype N2 *C. elegans*. Quantitative LA-ICP-TOF MS ion images of endogenous elements in day 1 gravid adult wildtype N2 *C. elegans*. Multi-element overlay image of $^{56}$Fe (green), $^{55}$Mn (red), and $^{66}$Zn (blue) (G). Lower limits for all elements were set to 0 ppm and upper limits were set to 99$^{th}$ percentile, with the upper limit for $^{55}$Mn set to 99.9$^{th}$ percentile. Embedded petrolatum shows similar localization of all elements as petrolatum-coated worms.

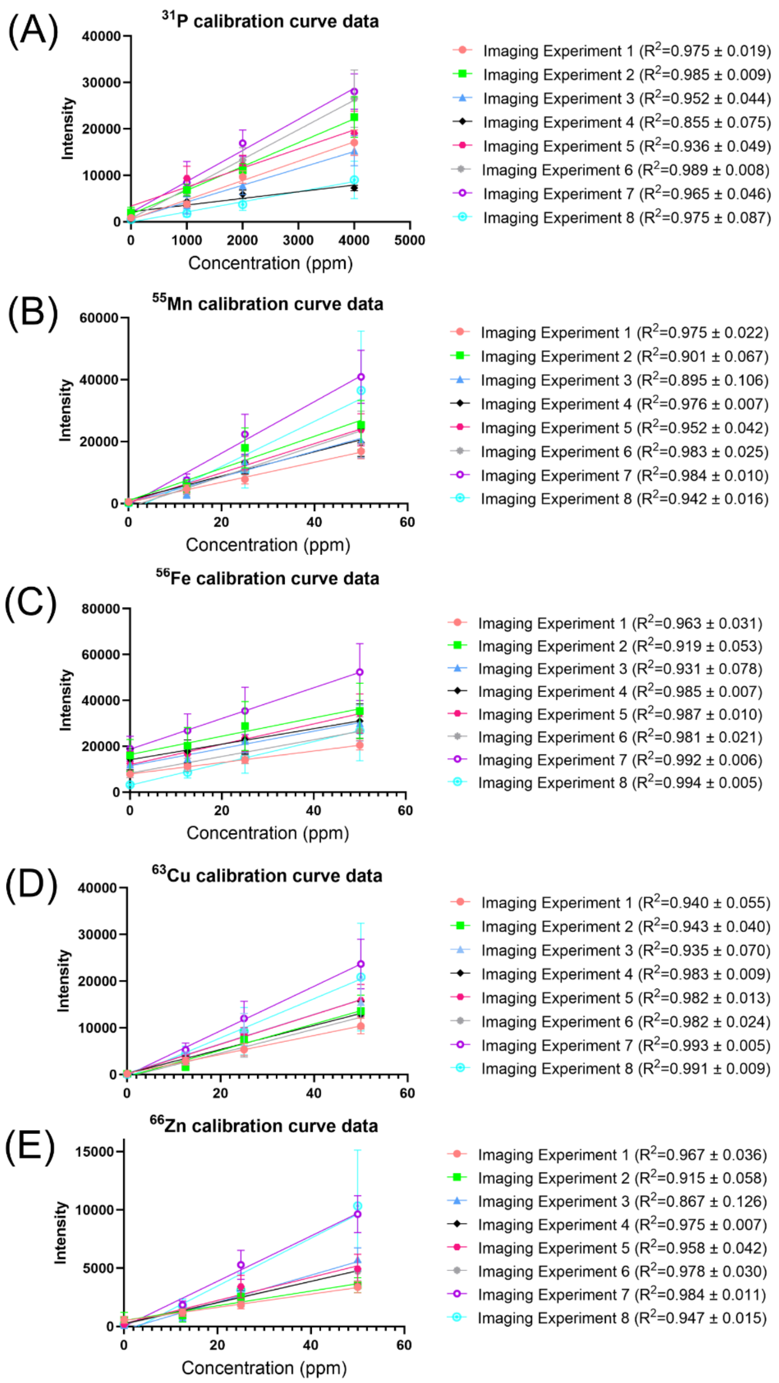


**Figure S13:** Raw calibration curve data for all quantitative LA-ICP-TOF-MS imaging of *C. elegans.* (A) $^{31}$P , (B) $^{55}$Mn, (C) $^{56}$Fe, (D) $^{63}$Cu, (E) and $^{66}$Zn calibration curves visualizing standard curves from multi-day imaging experiments. $R^2$ averages and standard deviations were calculated from the $R^2$ values of repeated gelatin scan measurements throughout individual runs. All imaging runs were performed on different days to assess inter-day variance. Data not background subtracted and forced through origin (0,0) to visualize blank intensity drift for all measured elements.

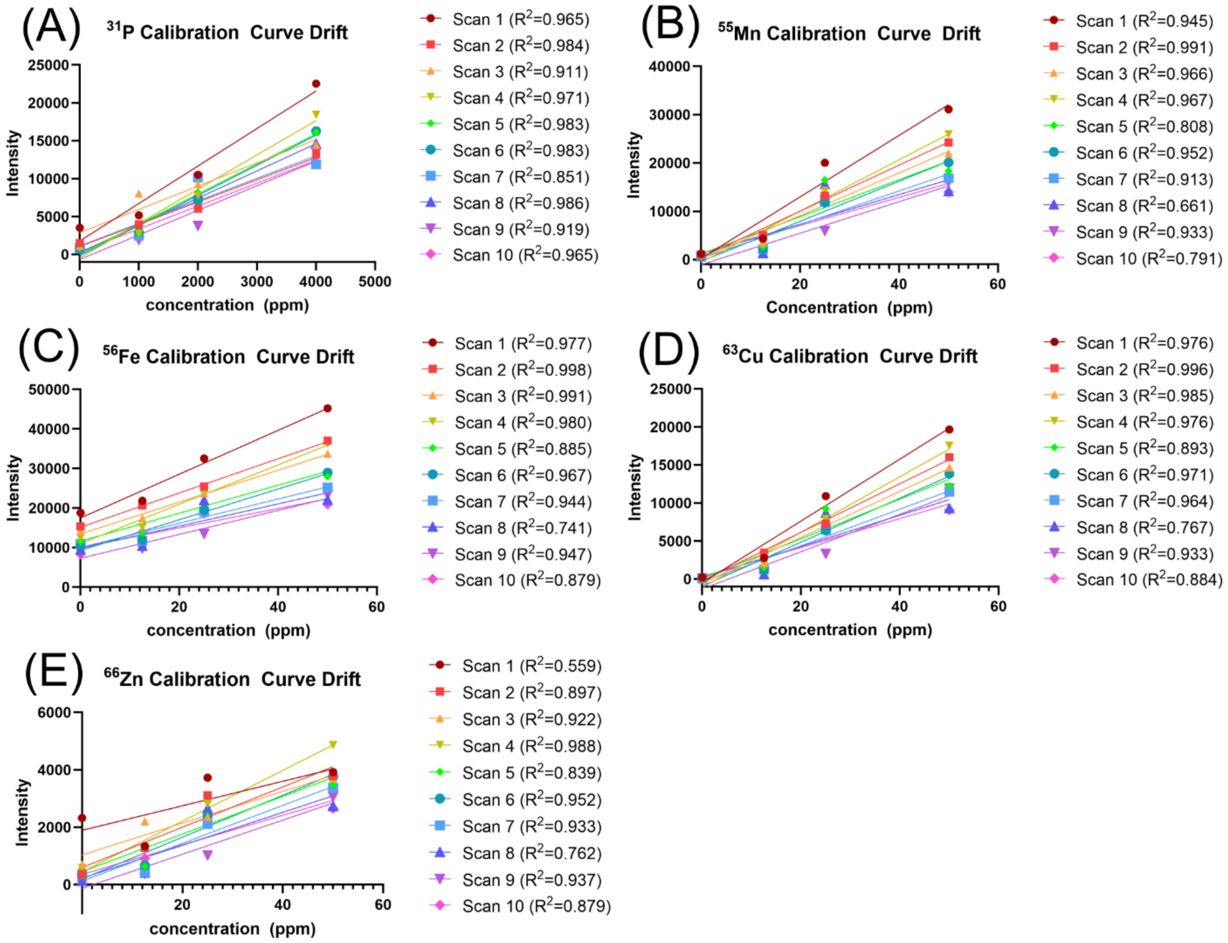


**Figure S14:** Calibration standard scan lines during a single LA-ICP-TOF-MS imaging experiment showing calibration sensitivity drift for all elements. (A) $^{31}$P, (B) $^{55}$Mn, (C) $^{56}$Fe, (D) $^{63}$Cu, (E) and $^{66}$Zn show both intensity and sensitivity drift throughout an imaging run. Instrument sensitivity drift contributes to inflated relative standard deviations of pooled calibration standard replicates. Data not background subtracted and forced through origin (0,0) to visualize blank intensity drift for all measured elements.

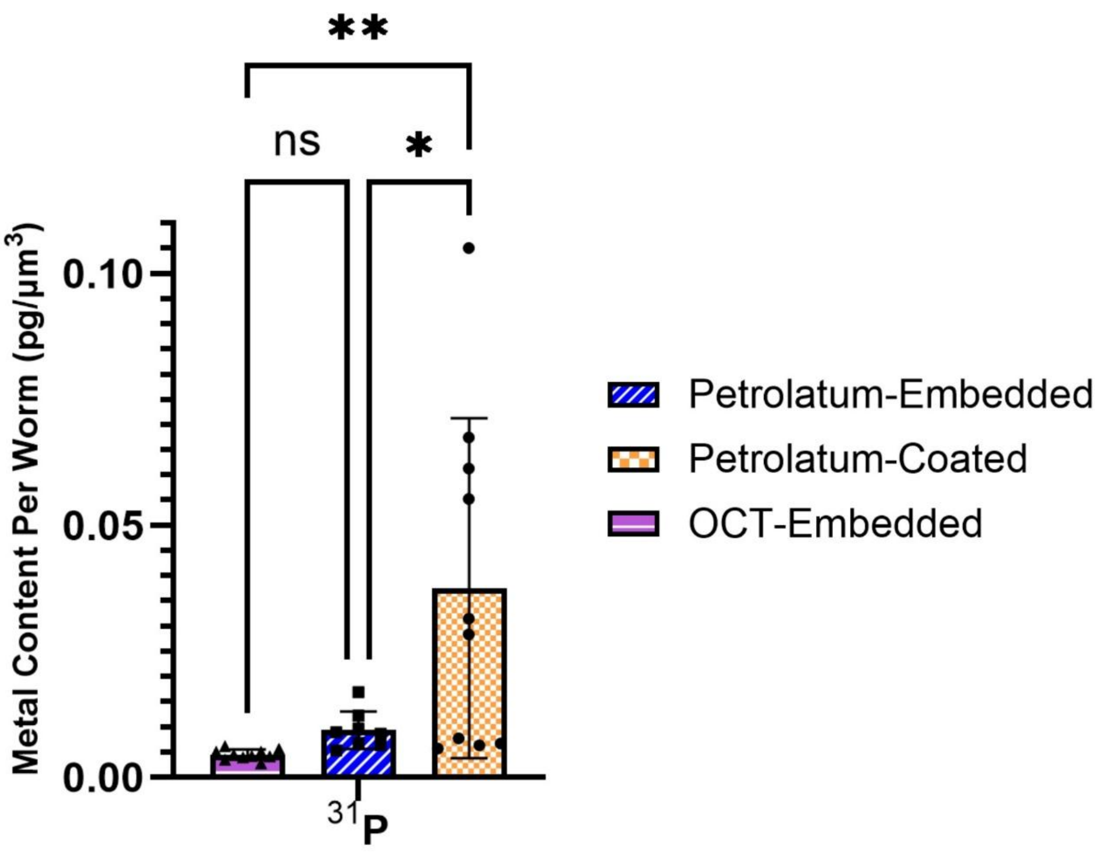


**Figure S15:** $^{31}P$ content normalized to individual *C. elegans* volume. $^{31}P$ content was measured to evaluate if worms were sufficiently rinsed of phosphate content present in M9 buffer during collection. OCT-embedded and petrolatum-embedded groups were found to be sufficiently cleaned of residual M9 buffer, yielding statistically similar content. 6 worms from the petrolatum-coated showed high levels of $^{31}P$ indicative of residual phosphate-containing M9 buffer on the surface of their bodies. *= $p<0.05$, **=$p<0.01$, One-way ANOVA; Tukey's HSD.

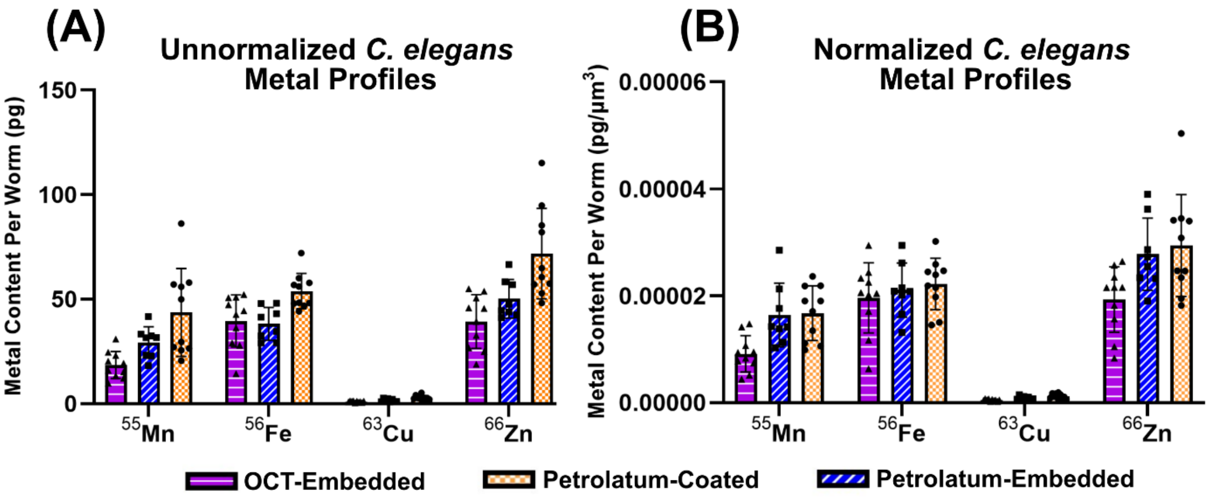


**Figure S16:** Total metal content determination in *C. elegans* by quantitative LA-ICP-TOF-MS imaging prepared using different sample preparation techniques. (A) Unnormalized metal content from *C. elegans* reported as picograms (pg) show large differences between all three sample preparation techniques. (B) Metal content normalized to the imaged worm's calculated volume (pg/µm$^3$) reduces metal content differences due to developmental variation.

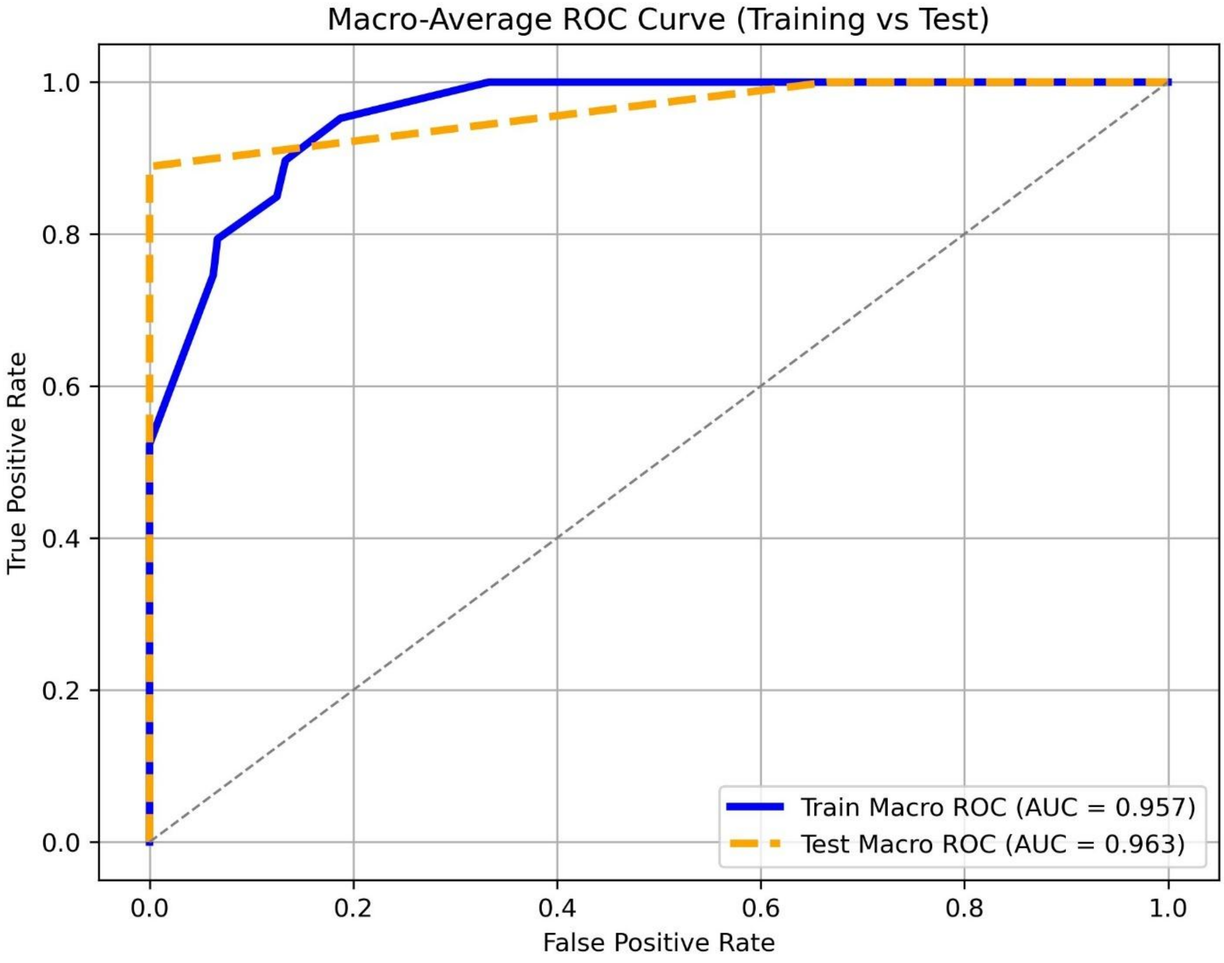


**Figure S17:** Evaluation of LDA model performance for class separation of OCT-embedded, petrolatum-embedded, and petrolatum-coated *C. elegans* used in post-hoc testing of MANOVA data. Both training and test sets yielded AUC values above 0.95 suggesting the model accurately separated classes. Training split used 80% of data (22 randomly selected worms) and test split used 20% of data (6 randomly selected worms).
